# Active Translation Control of CD4 T Cell Activation by Regulatory T Cells

**DOI:** 10.1101/2021.09.23.461566

**Authors:** Lomon So, Kazushige Obata-Ninomiya, Alex Hu, Virginia Muir, Ayako Takamori, Jing Song, Jane H. Buckner, Ram Savan, Steven F. Ziegler

## Abstract

Increased protein synthesis is a hallmark of lymphocyte activation. Regulatory T cells (Tregs) suppress the activation and subsequent effector functions of CD4 effector T cells (Teffs). However, molecular mechanisms that enforce Treg-mediated suppression in CD4 Teff are not fully clear. Control of CD4 Teff activation by Tregs has largely been defined at the transcriptional level, which does not reflect changes in post-transcriptional control. We found that Tregs suppressed activation-induced global protein synthesis in CD4 Teffs prior to cell division. We analyzed genome-wide changes in the transcriptome and translatome of activated CD4 Teffs using two independent approaches. We show that mRNAs encoding for the protein synthesis machinery are regulated at the level of translation in activated Teffs. Strikingly, Tregs suppressed global protein synthesis of CD4 Teffs by specifically inhibiting mRNAs of the translation machinery at the level of mTORC1-mediated translation control. Lastly, we found that the RNA helicase eIF4A inhibitor rocaglamide A (RocA) can suppress CD4 Teff activation *in vitro* to alleviate inflammatory CD4 Teff activation caused by acute Treg depletion *in vivo*. These data provide evidence that peripheral tolerance is enforced by Tregs through mRNA translational control in CD4 Teffs. Therefore, therapeutic targeting of the protein synthesis machinery can be expected to mitigate inflammatory responses invoked by Treg loss of function.

## INTRODUCTION

Most self-reactive T cells are eliminated in the thymus through the process of central tolerance. However, a small percentage of cells escape to the periphery, where they have the potential to promote autoimmunity. These cells are normally held in check by a population of CD4 T cells referred to as regulatory T cells (Tregs). Tregs are essential to maintain immune homeostasis, and the transcription factor FOXP3 has been shown to be central to the development and function of Tregs. Mutations in the FOXP3 gene in mice and human patients with IPEX syndrome drive development of a common set of autoimmune symptoms(*1–4*). Mutations in the FOXP3 gene and the autoimmune phenotype are linked to a loss of Tregs or their function(*2, 5*). Tregs have the ability to potently suppress CD4 effector T cells (Teff) either directly or through the modulation of antigen presenting cells (mainly DCs) to ultimately suppress activation, proliferation, and subsequent effector functions of Teffs (*6–8*). Several mechanisms have been proposed for Treg-mediated suppression, including release of suppressive cytokines (e.g., TGFß, IL-10, IL-35) and expression of inhibitory receptors (e.g., CTLA-4, PD-1, TIGIT)(*9, 10*). Although a block in proliferation and effector T cell function have been the hallmarks of Treg mediated suppression, the molecular changes in target Teffs following Treg encounter remains unclear. This is especially true for the first 24 hours prior to the onset of Teff proliferation, when the biosynthetic capacity of the cell is greatly expanded(*11, 12*).

Upon activation, resting T cells undergo a rapid biosynthetic and metabolic reprogramming in preparation for cell division(*12–15*). Included in this reprogramming is an increase in translational activity and capacity(*16, 17*). In this study, we show that Tregs suppress activation of Teffs by enforcing a global inhibition of mRNA translation. We assessed the genome-wide changes in transcriptome and translatome in activated CD4 Teffs and identified translation control of mRNAs encoding components of the protein synthesis machinery. In the first 24 hours of Teff activation, a set of mRNAs encoding proteins involved in the translational machinery are shifted to polysomes, with no concomitant changes in their transcription. We found that Tregs specifically inhibit the shift of these mRNAs to polysomes by suppressing mammalian target of rapamycin complex 1 (mTORC1) signaling. In support of these findings, we provide new evidence that with direct targeting of protein synthesis using rocaglamide A (RocA), an RNA helicase eIF4A inhibitor, inflammatory CD4 Teff activation caused by *in vivo* Treg loss can be alleviated. In summary, we provide a novel mechanism of Treg-mediated suppression of CD4 Teff activation through inhibition of mRNAs encoding protein synthesis machinery at the post-transcriptional level and that this biological mechanism can be therapeutically targeted using small molecule inhibitors.

## RESULTS

### Tregs control the protein synthetic capacity of activated CD4 Teffs

The 24-48 hours following CD4 Teff activation are critical for subsequent proliferation and expansion. This is a period during which cellular biomass is accumulated through expansion of global protein synthetic capacity in preparation for cell division. We reasoned that this period could be a target for Treg-mediated suppression in order to inhibit cell activation prior to proliferation. Overall protein synthesis rate can be quantified at the single-cell level by pulsing cells with the tRNA-analog puromycin (PMY) and intracellular staining for PMY. We co-cultured CD4 Tconv cells (CD4^+^CD25^-^) and congenically marked Tregs (CD4^+^Foxp3^+^) at varying ratios with anti-CD3/CD28 coated beads and pulsed the culture with PMY. CD4 Tconv cells co-cultured with Tregs exhibited marked inhibition of proliferation in a dose-dependent manner (Figure S1A). Interestingly, Tconv cells co-cultured with Tregs clearly showed significantly less PMY incorporation in a Treg-dose-dependent manner before the onset of proliferation (Figure 1A). The pattern of suppression was not bimodal, indicating that the protein synthesis rate of all responding Tconv cells was modulated by Tregs, less completely than upon cycloheximide (CHX) treatment (Figure 1B). Since anti-CD3/CD28 beads were used to activate both populations of cells in the co-culture system, the downregulation of protein synthesis by Tregs is independent of antigen-presenting cells (APC). The Treg-mediated translational inhibition was observed as early as 6h post-activation, well before any metabolic changes occur in T cells (Figure 1C). Suppression was primarily due to early prevention of T cell activation as we found no difference in the ability of CD4 Teffs to downregulate CD62L with or without Tregs (Figure S1B). Thus, suppression of global protein synthesis in CD4 Teffs by Tregs could not be attributed to dampening or cold inhibition of general T cell activation. To test whether similar regulatory pathways were operative in human T cells, we established *ex vivo* Treg suppression assays using PMY incorporation as the readout. We used *in vitro-expanded* Tregs from a single donor, and Tconv (defined as CD4^+^CD45RA^+^CD127^+^CD25^-^) from 5 individual healthy donors. We found that 24h of stimulation resulted in a significant increase in PMY staining, both in percentage of cells labeled and the MFI of PMY staining. In the cultures containing Tregs, PMY incorporation was significantly reduced, both in total incorporation and in the percentage of cells that incorporated PMY (Figure 1D). These data demonstrate that Treg-mediated inhibition of activation-induced translation is a conserved function.

**Figure 1.**
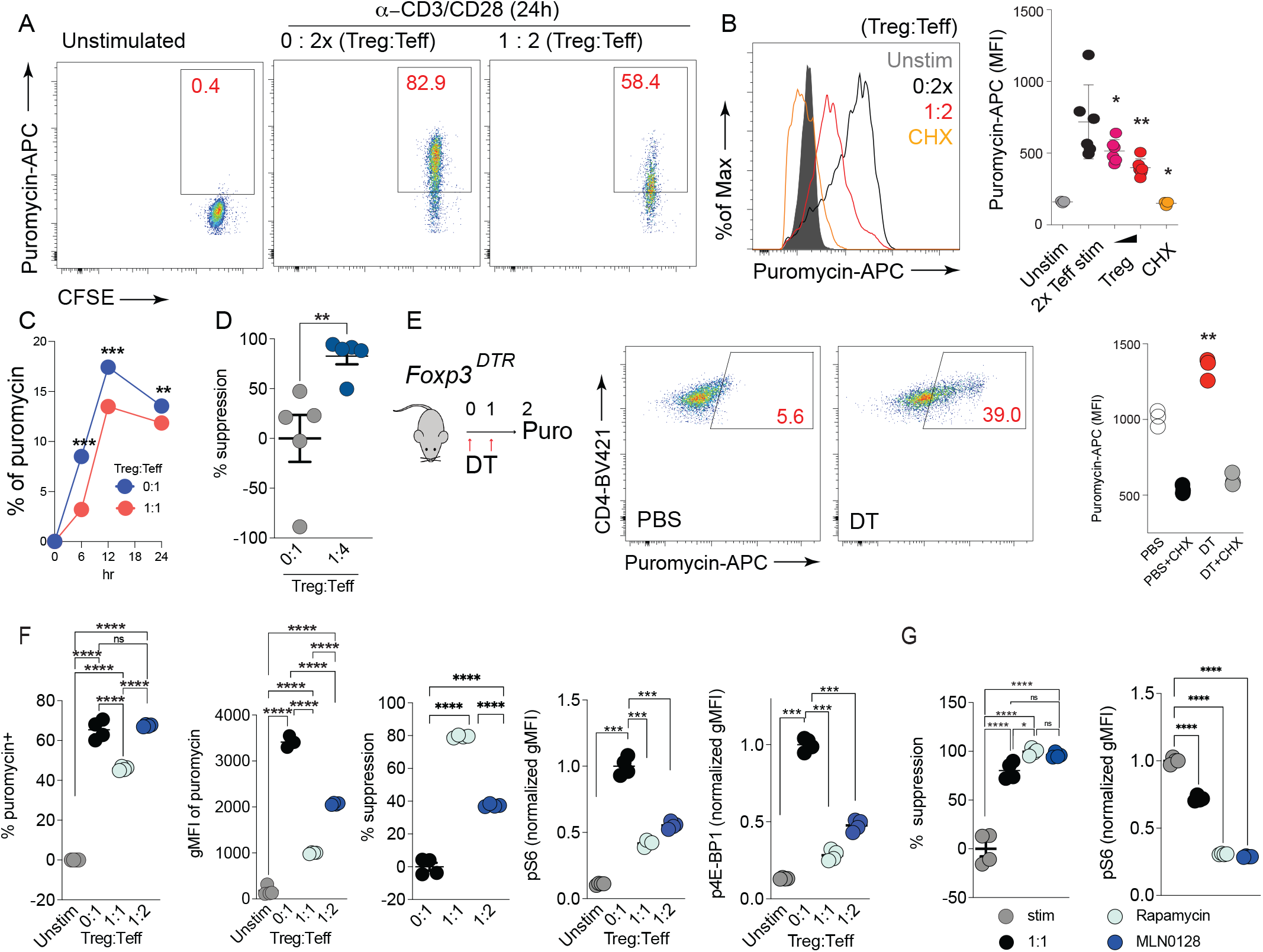
Tregs suppress global protein synthesis in CD4 Teffs. (A) Purified naïve CD4 T cells from *Foxp3^YFP-Cre^* mice (CD4+YFP-) were stimulated with equal number of anti-CD3/CD28 beads with or without the indicated ratio of Foxp3^+^Tregs (CD4+YFP+; Treg:Teff = 1:2). For cultures with CD4 Teffs only, cell number was doubled to match total cell number in the culture (2x). Cells were pulsed with puromycin (PMY) for 15min before harvest and intracellular staining for incorporated puromycin performed using a fluorophore-conjugated anti-PMY antibody. (B) Purified naïve CD4 T cells were stimulated with anti-CD3/CD28 beads with or without FACS sorted Foxp3-YFP+ Tregs from *Foxp3^YFP-Cre^* mice (Treg:Teff = 1:2) for 24h, followed by analysis of PMY incorporation as in (A). PMY signal and median fluorescence intensity (MFI) were quantified. Cycloheximide (CHX) was added to 24h stimulated CD4 Teffs 5min prior to PMY pulse to act as a negative control and set the baseline protein synthesis signal. One-way ANOVA was applied. (C) Purified naïve CD4 T cells from *Foxp3^YFP-Cre^* mice as CD4 Teffs were stimulated with anti-CD3/CD28 beads with or without the indicated ratio of FACS sorted Foxp3^−^YFP^+^ Tregs from *Foxp3^YFP-Cre^* mice (Treg:Teff = 1:1) for 6, 12, 24h, followed by analysis of PMY incorporation as in (A). (D) *In vitro*-expanded Tregs from a single donor, and Teff (defined as CD4^+^CD25^-^CD45RA^+^) from 5 individual healthy donors were used for the human Treg suppression assay. Cells were co-cultured and stimulated with anti-CD3/CD28 beads for 24h and PMY staining was performed as in (A). The % suppression was calculated as described in materials and methods. (E) Foxp3^DTR^ mice were injected with diphtheria toxin (DT) for two consecutive days and splenocytes were harvested on day 2. Protein synthesis in CD4 T cells was measured by *ex vivo* puromycin pulsing of splenocytes and gating on CD4 T cells. (F) CD4 Teffs stimulated in the absence and presence of Tregs were identified by congenic markers and intracellular signaling molecules were assessed by phospho-flow cytometry. (G) Purified naïve CD4 T cells were treated with 5 nM Rapamycin and 50 nM MLN0128 for 30 min prior to stimulation for indicated samples. The cells were harvested with or without equal number of anti-CD3/CD28 beads in the presence or absence of iTregs (Treg:Teff = 1:1) for 24 h, subjected for analyzing PMY incorporation and phosphorylation of S6 by flowcytometry. One-way ANOVA was applied for all comparisons. \**P*<0.05, \*\**P*<0.01, \*\*\**P*<0.001. All experiments were repeated at least twice.

Next, to assess the role of Tregs in controlling protein synthesis in CD4 T cells *in vivo*, we acutely depleted Tregs through diphtheria toxin (DT) treatment of Foxp3^DTR^ mice (*18*). Within 3 days post initial DT-induced depletion of Tregs (2 consecutive DT injections on days 0 and 1), we observed rapid appearance of a CD4 T cell population with significantly elevated incorporation of puromycin *ex vivo* compared to the PBS treated control mice, suggesting activation of the autoreactive CD4 T cell pool in the periphery (Figure 1E). When CD4 T cells from spleen and lymph nodes of DT-treated Foxp3^DTR^ mice were purified and stimulated *ex vivo*, they proliferated with faster kinetics and were significantly larger in size, indicating an aberrantly enhanced protein synthesis capacity in Teffs activated *in vivo* attributed to Treg loss (Figure S1C). These data suggest that Tregs are both sufficient and necessary to suppress the rapid upregulation of protein synthesis in activated CD4 T cells both *in vitro* and *in vivo*.

To uncover the underlying mechanism of Treg-mediated translational inhibition, we examined signaling pathways downstream of TCR stimulation in activated CD4 T cells. Specifically, we examined the mammalian target of rapamycin (mTOR) signaling pathway as it has been shown to be critical for coordinating cell growth and proliferation in lymphocytes through the eukaryotic translation initiation factor 4E (eIF4E) in translation initiation (*19*). mTOR exists in 2 multi-protein complexes, mTOR complex 1 (mTORC1) and mTOR complex 2 (mTORC2) (*20, 21*). To assess mTORC1 signaling, we examined phosphorylation of ribosomal protein S6 (rpS6: S240/244) and eIF4E-binding proteins (p4EBP1/2: T37/46) at their respective mTORC1 specific phosphosites in activated CD4 T cells. mTORC2 signaling was assessed by phosphorylation of AKT at S473. As expected, both mTORC1 and mTORC2 signaling increased in activated CD4 T cells. Strikingly, Tregs significantly suppressed mTORC1 signaling (rpS6 and p4EBP1/2) (Figure 1F). However, mTORC2 signaling (AKT S473) remained intact, as did the PDK1- and PI3K-dependent phosphosite T308, which is more proximal to TCR engagement (Figure S1D). Using Nur77-GFP reporter Teffs, we see no differences in Teffs alone or co-cultured with Tregs, indicating no change in proximal TCR signaling (Figure S1E). Furthermore, treatment of Teffs with mTORC inhibitors Rapamycin or MLN0128, suppressed phosphorylation of rpS6 and puromycin incorporation to the same extent as Treg-mediated suppression (Figure 1G). These data suggest that Treg-mediated translational inhibition in Teffs is associated with reduction of mTORC1 signaling in Teffs. These findings are consistent with previous data showing that genetic or chemical inhibition of mTORC1 significantly inhibits lymphocyte proliferation in a 4EBP/eIF4E-dependent manner to control translation initiation in various cell types, including lymphocytes (*11, 22*).

### Development of a Simple Polysome Efficient Extraction and Distribution (SPEED) technique to identify mRNAs that are differentially translated

The transition of mRNAs from monosome to polysome and back is a critical aspect of translation control. To resolve and distinguish mRNAs bound to polysomes from monosome-associated mRNAs, we optimized the classical polysome profiling approach for low input cytosolic lysates (Figure 2A). We reasoned that by assessing the quantity and quality of total RNA extracted from each fraction, we could determine ribosome positions since total RNA is mainly composed of ribosomal RNA (rRNA). Using lysates prepared from as few as 500,000 to 1 million activated CD4 T cells, we found that total RNA extracted from each fraction and plotted as a percent distribution plot closely resembled a classical A254nm polysome trace obtained using >20 million activated CD4 T cells (Figure 2B). We have termed this a ‘<u>S</u>imple <u>P</u>olysome <u>E</u>fficient <u>E</u>xtraction and <u>D</u>istribution’ (SPEED) plot to distinguish it from the traditional polysome traces requiring greater cellular input (Figure 2C). Furthermore, qualitative analysis of the extracted RNA using Bioanalyzer gave us information as to the position of the intact 80S monosome in the gradient (Figure 3C). The SPEED plots generated from unstimulated CD4 T cells showed enrichment of most of the total RNA in the monosome fraction (Figure 2C), confirming our observations that CD4 T cells prior to activation have low translational activity (Figure 1A). Upon stimulation, nearly 50% of the monosomes shifted towards the heavier sucrose fractions, indicating the assembly of polysomes and increased protein synthesis (Figure 2C). To ensure SPEED plots faithfully represent the mRNA translational status of a cell, we took advantage of the translation initiation inhibitor homoharringtonine, which only interferes with initiating ribosomes and allows elongating ribosomes to run-off(*23*). Activated CD4 Teffs were treated with homoharringtonine (HHT) for 10 min to allow run-off elongation of ribosomes before cytosolic lysate preparation. Remarkably, the SPEED plot from HHT treated activated CD4 Teffs resembled unstimulated CD4 T cells, with majority of ribosomes enriched in the monosome fraction (Figure 2C). Lastly, the distribution of beta-actin (*Actb*) mRNA was analyzed using quantitative PCR (qPCR) from each fraction. We chose *Actb* as it is routinely used as a housekeeping control mRNA as it has no apparent *cis*-regulatory sequence motif in its 5’ untranslated region (UTR) and is highly translated. Despite low polysome levels in unstimulated CD4 Teffs, *Actb* mRNA was abundantly enriched in the polysome fractions. Activated CD4 Teffs also translated *Actb* mRNA with high efficiency. As expected for a highly translated mRNA, HHT treatment led to a complete shift in *Actb* mRNA towards the lighter sucrose fractions, indicating successful ribosome run-off (Figure 2D). In summary, our SPEED technique faithfully captured the translational status of cellular lysates from low biological input, making it ideal to assess the translatome of primary immune cells, and most importantly, of Treg suppressed CD4 Teffs.

**Figure 2.**
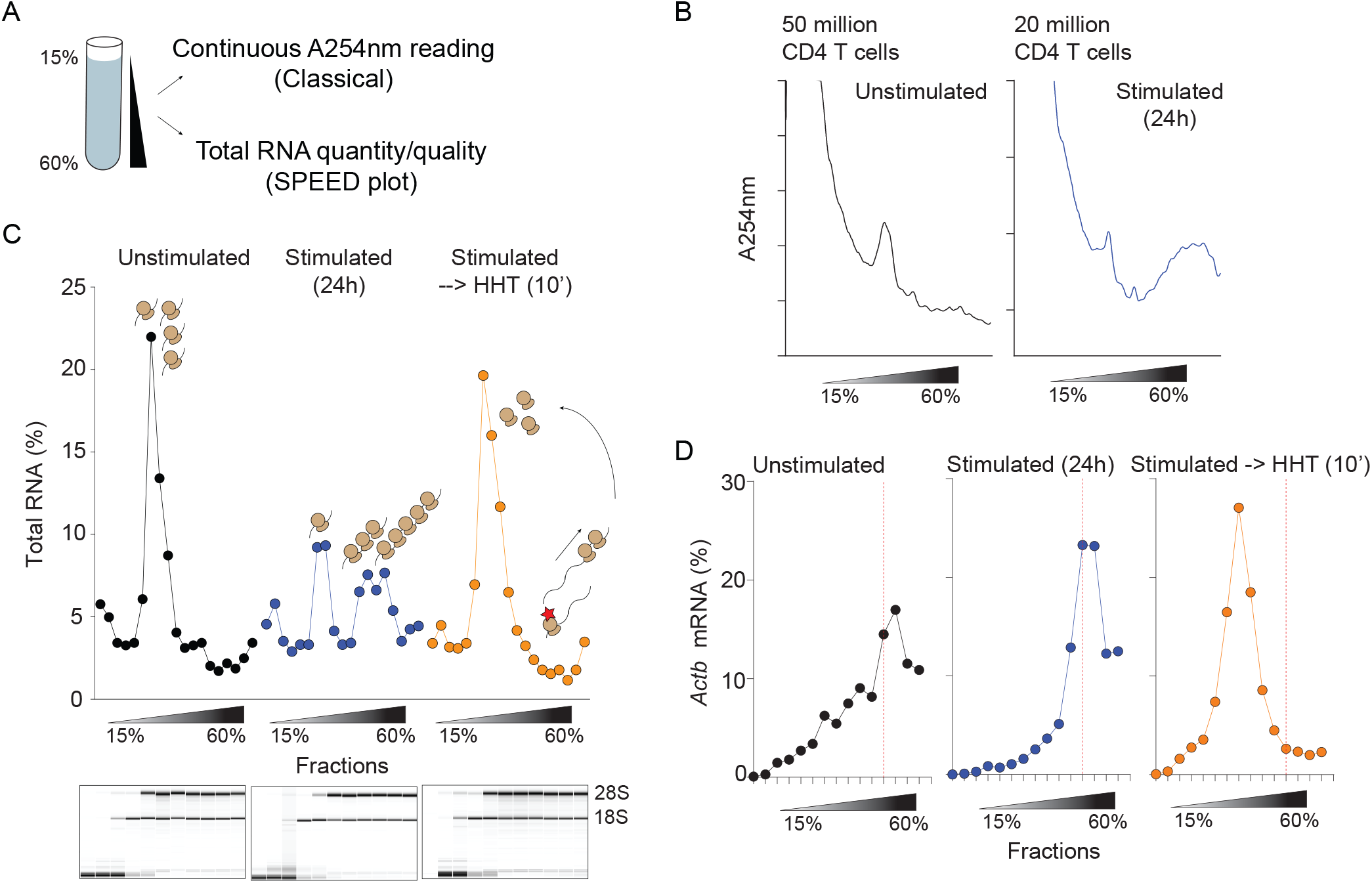
SPEED analysis as a novel polysome profiling approach. (A) Polysome profiling approach using sucrose gradients to physically stratify cellular cytosolic lysates. Classically, continuous absorbance at 254nm (A254nm) is used to assess monosome and polysome positions. SPEED utilizes analysis of the quantity and quality of total RNA from each fraction, making it amenable for ultra-low biological input that is below the detection limit of A254nm reading. (B) Indicated numbers of bulk CD4 T cells were left unstimulated or stimulated with equal number of anti-CD3/CD28 beads for 24h and subjected for classical polysome profiling using A254nm reading to obtain polysome traces. (C) Same samples from (B) but equivalent to 500,000-1 million activated CD4 T cells and 2-3 million unstimulated CD4 T cells were subjected for polysome fractionation. No A254nm traces were obtained. Total RNA was extracted from each sucrose fractions and quantified to plot total RNA percent distribution across fractions. Total RNA quality was assessed using Bioanalyzer (bottom Bioanalyzer results; only fractions #1-11 were analyzed since each RNA pico chip can analyze 11 samples at a time). (D) Equal volume of RNA from each fraction was reverse transcribed into cDNA and *Actb* mRNA levels were quantified using qPCR. The percent *Actb* mRNA across fractions was quantified and plotted. Red dashed line indicates fraction #12 in each sample.

### Genes controlling mRNA translation are affected in T cell activation and Treg mediated suppression

To identify the mRNAs that shift between monosomes and polysomes following activation, we performed SPEED on CD4 T cells activated for 24h with anti-CD3/CD28 beads. A portion (10%) of cellular lysate was used to isolate total RNA, while the remainder was subjected to SPEED analysis with monosome and polysome fractions collected and RNA isolated and sequenced (Figures S2 and S3). We found approximately 348 genes with differential translational efficiency (Translation efficiency (TE) was calculated as polysomal/subpolysomal enrichment) when compared stimulated versus unstimulated cells, with 199 and 149 genes with lower and higher TE, respectively (Figure 3B). Among them, 201 genes changed in both TE and overall RNA expression in the same direction, 97 genes change in just TE and not overall input RNA and 50 gene changed in the opposite direction (Figures 3C-D, S3A-B, supplement table 1). Gene ontology analysis of differential TE genes showed that the vast majority fell into functional categories involving ribosome biogenesis and mRNA translation, consistent with preparation for subsequent cell division following activation. It is important to note that the genes that encode these mRNAs are considered to be housekeeping genes whose expression is very high and largely unchanged by cell activation. The data presented here demonstrates that while the RNA level of transcripts encoding the transcriptional machinery is unchanged, their translation is likely increased following activation due to their shift onto polyribosomes. We next determined the fate of these mRNAs in Teffs stimulated in the presence of Tregs. We examined the mRNAs that showed increased translation efficiency (TE) in stimulated vs. resting cells and calculated their TE in CD4 T cells stimulated in the presence of Tregs (Stim+Treg). The set of mRNAs showing increased TE in stimulated cells were specifically reduced in their TE when the cells were stimulated in the presence of Tregs (Figure 3E), even though the total mRNA levels of these genes remain unchanged (Figure S3C). These data are consistent with our hypothesis that Treg-mediated translational control targets mRNAs involved in preparing the cell for subsequent proliferation, with ribosome biogenesis and translational activity being critically important for this process. The finding that stimulation in the presence of Tregs reversed the positive TE changes in transcripts encoding proteins involved in mRNA translation is consistent with data showing that overall protein synthesis is suppressed by Tregs (Figure 3F).

**Figure 3.**
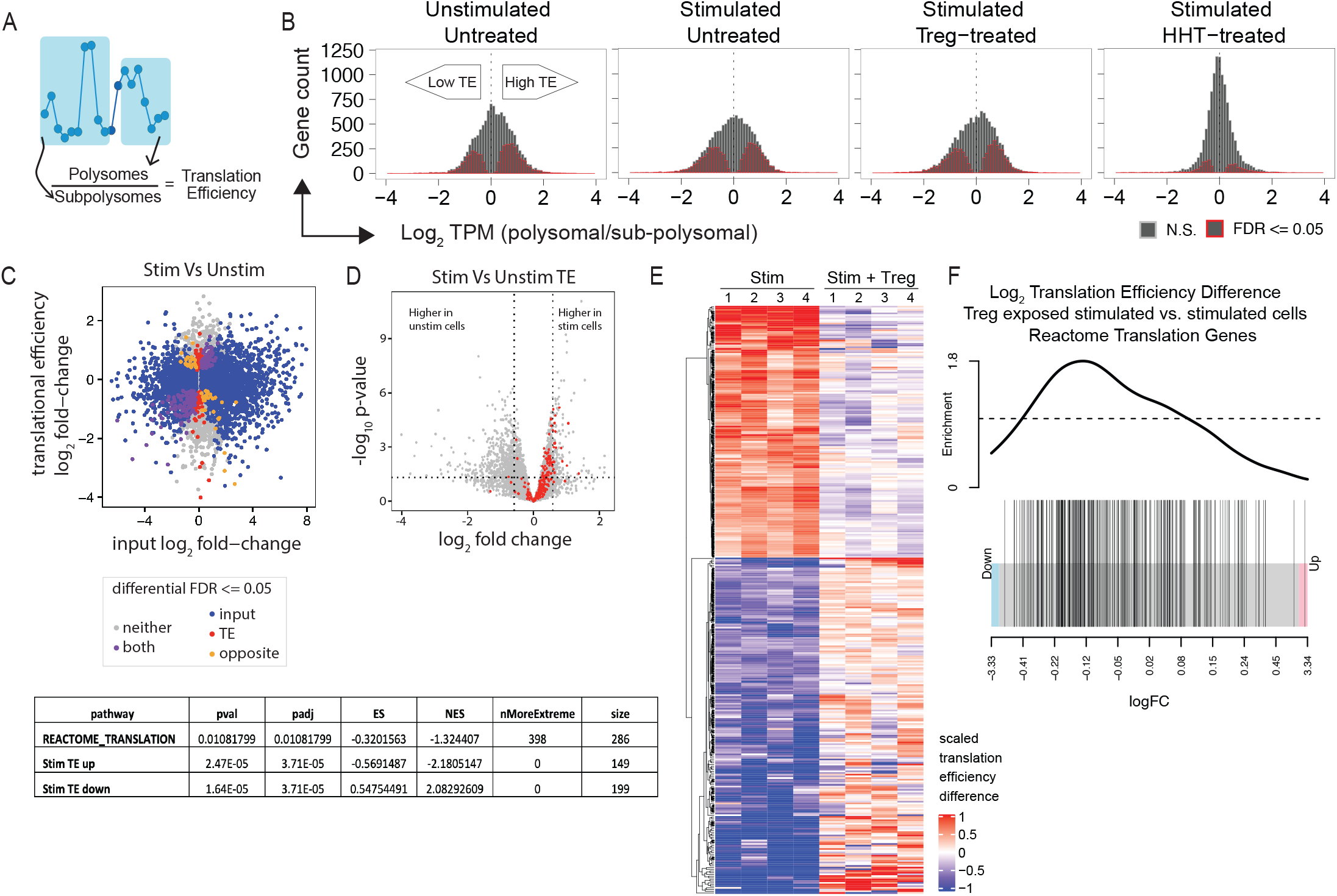
Tregs specifically suppress mRNAs related to the translational machinery via active translation control. (A) Scheme to capture polysome and subpolysome-associated mRNAs for next-generation sequencing. (B) Translation efficiency (TE) was calculated as polysomal/subpolysomal enrichment for four conditions; 1) Unstimulated CD4 Teffs, 2) 24h stimulated CD4 Teffs, 3) 24h stimulated CD4 Teffs co-cultured with Tregs, 4) 24h stimulated CD4 Teffs treated with harringtonine for 30min. Histograms of log_2_ translation efficiency (TE) for all genes were computed for the four conditions. Genes whose log_2_ TE were different than 0 at £ 5% false-discovery rate (FDR) are highlighted in red. (C) Log_2_ fold-change of gene expression in the input RNA is plotted against log_2_ fold-change of translational efficiency between the stimulated and unstimulated cells. The colors annotate genes based on the significance of their input differential expression or translational efficiency-red if the gene changes in translational efficiency but not expression, blue if the gene changes in expression but not translational efficiency, purple if the gene changes in both, grey if the gene changes in neither, and orange if the gene changes in both but in opposite directions. (D) Volcano plot showing log fold-changes of TE in genes between stimulated and unstimulated CD4 Teffs and their adjusted p-values. Dots in red are genes of the Reactome translation pathway. (E) Heatmap of log fold-change of TE in stimulated CD4 Teffs and stimulated CD4 Teffs co-cultured with Tregs, both against unstimulated CD4 Teffs. (F) Barcode plot ranks genes by their log fold-change of TE between Treg-exposed stimulated cells and stimulated cells (p-value 0.009377618). Genes highlighted in black are genes within the Reactome Translation gene set identified to have increased TE in the stimulated vs. unstimulated cell comparison. P-value are computed using the fgsea (62) package that implements Gene Set Enrichment Analysis (GSEA) statistics.

### mRNAs sensitive to translational control in CD4 T cells are enriched for the terminal oligopyrimidine (TOP) motif

To identify the mechanism(s) of translation control, we first examined *cis*-regulatory elements shared by the 5’UTRs with higher TEs identified by SPEED (Supp material 2A-E). We searched for sequence-specific motifs within target UTRs in these Treg sensitive genes by retrieving murine 5’UTR sequences from the Ensembl BioMart database (Ensembl Genes78, Mus musculus) and using Multiple Em for Motif Elicitation (MEME), to uncover common *cis*-regulatory elements(*24–26*). Briefly, 131 genes increased in TE in stimulated cells relative to unstimulated cells at 5% FDR and also decreased in TE in Treg-exposed stimulated cells relative to the unstimulated cells (2A). Of those 131, we were able to extract 5’ untranslated mRNA sequences for 127 sequences (2B) using biomart (http://uswest.ensembl.org/biomart/martview/e728a9d413ab0dfd039a3ce666b3d3e2). We used the MEME software tool to identify de novo a motif enriched in these sequences (considering both the positive and negative strand), and the top hit closely resembles a TOP motif (2C). 109 of the 127 sequences have an instance of the motif with a p-value of 0.01 or lower. 82 of the 109 show the motif in the strand orientation we expect, while 27 show the motif in the opposite orientation. To confirm that this motif is still significantly enriched in the correct orientation, the motif was tested for enrichment using the SEA tool, also in the MEME Suite package, on only the correct orientation. The motif was enriched at a p-value of 0.000126 (2D), and this tool identified 67 instances of the motif among the 127 sequences (2E). We found an oligopyrimidine tract enriched in the 5’UTR in Treg sensitive mRNAs (Figure 4A). The terminal oligopyrimidine (TOP) motif is a well-characterized motif which regulates key mRNAs encoding the translational machinery. The TOP motif is known to be present in the 5’UTR of ribosomal proteins (RPs) and other genes required for mRNA translation(*27*). Interestingly, TOP motif containing RP transcripts are regulated by the well-known signaling kinase mTORC1(*22, 28*). To validate select genes that were differentially regulated in SPEED RNA-seq. We observed that *Rps10, Rpl14, eIF3e*, and *Rpl8* mRNAs that contain TOP motif (Figure SF4A), that are involved in mRNA translation, shift to polysomes upon CD4 T cell stimulation and back to monosomes following activation in the presence of Tregs (Figures 4B, D). The translational block in Treg coculture does not reflect a change in the total RNA levels of these genes (Figure 4C), but rather a change in the translation of these mRNAs. As predicted, the mRNA distribution shifted back to the monosome fraction when the CD4 T cells were stimulated in the presence of Tregs. As a control, *ActB* mRNA distribution is unchanged by Tregs. Finally, we also observed changes of TE of Rpl8 and eIF3e also reflected at the protein levels measured by flow-cytometry (Figures 4E, SF4B-E). These data confirm global changes in the translatome induced by Tregs to control CD4 T cell activation by blocking the translation of a subset of mRNAs, thereby blunting the ability of these cells to respond appropriately to stimulation.

**Figure 4.**
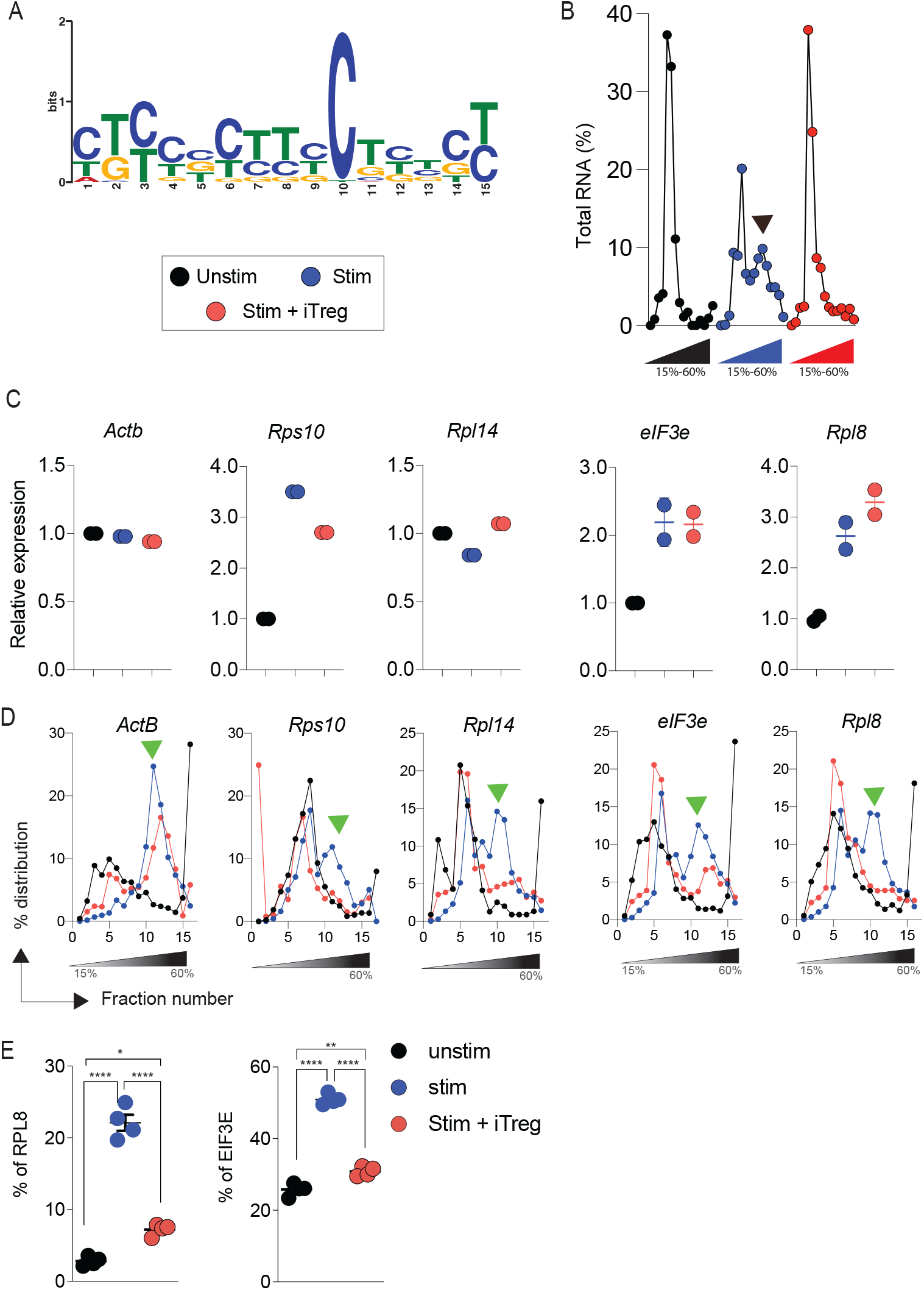
Tregs suppress translation of TOP motif containing mRNAs. (A) Motif discovery analysis finds a TOP motif in Treg-regulated mRNAs. (B) SPEED profiling of unstim, stim and stim+iTreg cells. Total RNA of low-input samples measured at A254nm. (C-D) mRNAs regulated by Tregs in Fig. 3 were validated using SPEED polysome qPCR. pPCR of RNA from input (C) and SPEED-fractionated (D) samples. Actb mRNA served as a control. The above experiments were repeated twice. (B,D) Green and black arrowheads highlight translation differences. (E) Purified naïve CD4 T cells were harvested with or without equal number of anti-CD3/CD28 beads in the presence or absence of equal number of iTregs for 6h, subjected for analyzing expression of protein levels of RPL8 and eIF3e by flowcytometry.

### IL-10 and TGFβ from Tregs block protein synthesis in Teffs

Since the activation of CD4 T cells in the presence of Tregs resulted in a specific inhibition of mTORC1 activation, as indicated by the lack of ribosomal protein S6 and 4E-BP1/2 phosphorylation, we hypothesized that Tregs suppress the immediate biosynthetic response to antigen-specific activation through regulation of mTORC1-mediated translational control. To define the pathways leading to the Treg-mediated inhibition of mTORC1 activation in stimulated CD4 T cells, we examined the role of soluble factors known to be produced by Tregs, using recombinant proteins, neutralizing antibodies, genetic knockouts specific for anti-inflammatory factors. It has been well known that deprivation of IL-2 is one of the mechanisms of Treg-mediated suppression of proliferation of Teffs. Consistent with this, when IL-2 signaling was blocked in stimulated T cells with αIL-2 or αCD25 neutralizing antibodies, we observed a reduction in pS6 and puromycin incorporation, while 4E-BP1 activation was not affected (Figure S5A).

We next examined whether IL-10 and TGFβ played a role in Treg-mediated translation inhibition. Blockade of IL-10 or TGFβ individually resulted in reduced inhibition of translation in Teff cells co-cultured with Tregs (Figure 5A). Importantly, neutralization of both IL-10 and TGFβ signaling significantly reduced Treg-mediated suppression of puromycin incorporation compared to neutralization of each signal respectively (Figure 5A). In addition, we found that blockade of IL-10 and TGFβ signaling lead to inhibition of Treg mediated suppression of mTORC1 signaling (Figure 5A). Similarly, we saw a significant rescue of protein synthesis and mTORC1 pathway when TGFβ was neutralized in IL-10Rb^-/-^ CD4 T cells co-cultured with Tregs (Figure S5B). These data suggest that Treg-mediated IL-10 and TGFβ, acting on Teff cells, result in a block in mTORC1 activation and subsequent translation inhibition. Consistent with this model we observed that addition of recombinant IL-10 and TGFβ to cultures of CD3+CD28-stimulated Teff cells resulted in significantly reduced puromycin incorporation and mTORC1 activation (Figure 5B). Finally, and consistent with these data, we have also found that Teffs activated in the presence of IL-10 and TGFβ resulted in a marked reduction of the polysome fraction, similar to what was seen in Teff-Treg co-cultures (Figures 4B and 5C). These data suggest that Tregs use production of IL-10 and TGFβ in combination to disrupt mTORC1 signaling and-inhibit mRNA translation in stimulated CD4 T cells.

**Figure 5.**
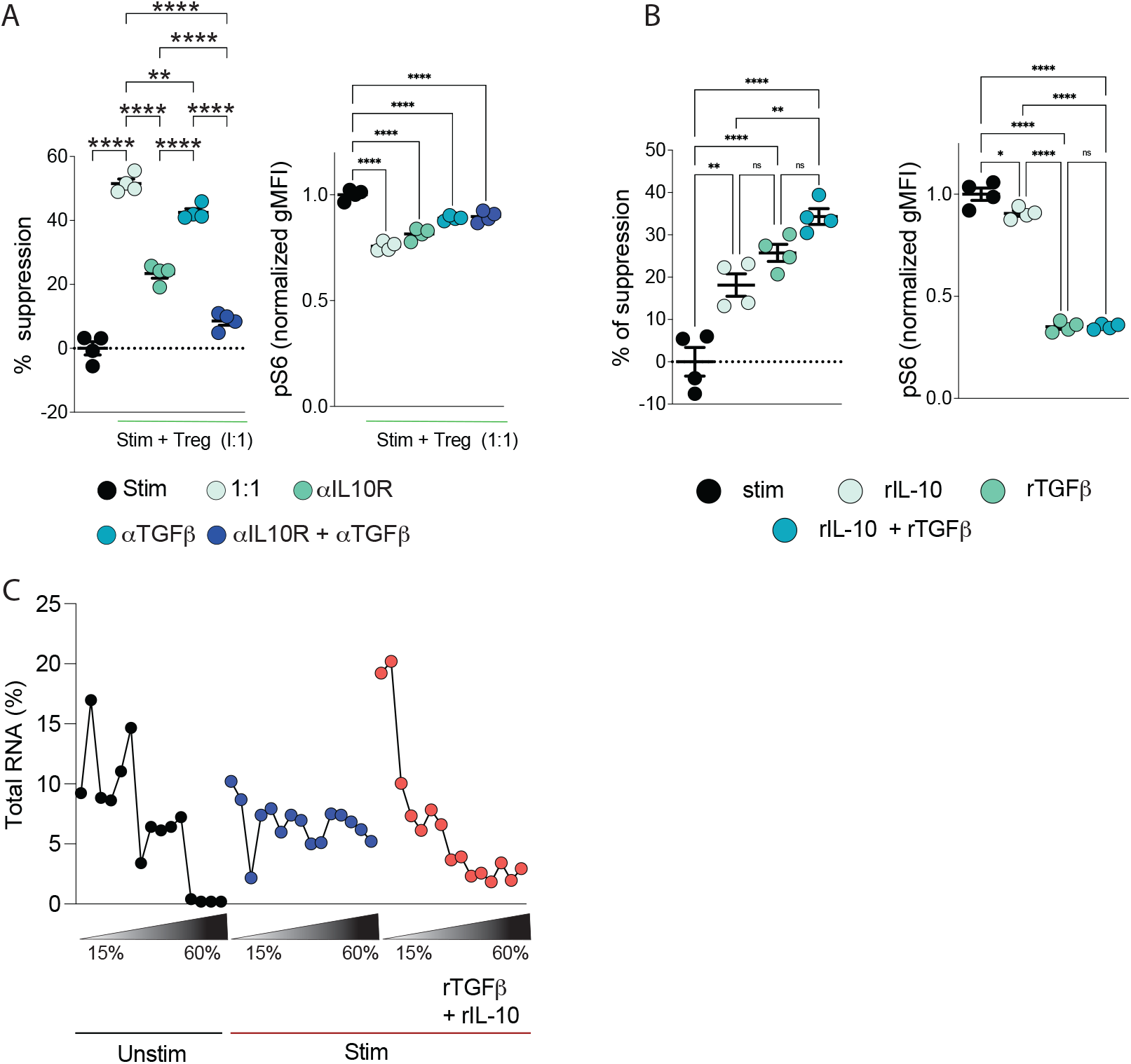
IL-10 and TGFβ from Tregs block protein synthesis in T effector cells. (A) Neutralizing antibodies for IL-10 receptor and/or TGFβ was added to co-culture of CD4 Teffs and Tregs with or without equal number of αCD3/CD28 beads for 24h, and analyzed for puromycin incorporation (left) and phosphorylation of S6 (right). (B) Recombinant IL-10 and TGFβ were added to culture of CD4 Teffs with or without equal number of anti-CD3/CD28 beads for 24h, and analyzed for puromycin incorporation (left) and phosphorylation of S6 (right). (C) SPEED profiling of unstim, stim and stim cells incubated with recombinant IL-10 and TGFβ. Total RNA of low-input samples measured at A254nm. (One-way ANOVA was applied for all comparisons. \**P*<0.05, \*\**P*<0.01, \*\*\**P*<0.001. The experiments were performed at least twice (A-C).

### mRNA translation inhibitor Rocaglamide A inhibits T cell proliferation

While we show that suppression of mRNA translation by Tregs has a direct effect on cell activation and proliferation, we next investigated if translation inhibition is one of the main ways Tregs exert their function downstream of mTORC1. To test this, we used a mRNA translation inhibitor Rocaglamide A (RocA,1*H*-2,3,3a,8b-tetrahyrocyclopenta[*b*]benzofuran), which has been shown to reduce overall protein synthesis but to preferentially inhibit specific subset of mRNAs in cell culture models (*29*). RocA is a secondary metabolite from the plant genus *Aglaia* with anti-tumor and anti-inflammatory properties (*30, 31*) (*30, 32, 33*). RocA affects protein synthesis by binding to eIF4A, a DEAD-box RNA helicase, that is part of the eIF4F mRNA initiation complex (*34*). RocA binds to eIF4A and clamps it on poly-purine sequences in the 5’UTRs of mRNAs, thereby preferentially inhibiting this set of mRNAs at the level of translation control when stable RocA-eIF4A-mRNA complexes are formed to block 43S pre-initiation complex scanning (*29, 35*). While RocA has been shown to inhibit overall protein synthesis through the translational blockade of a subset of mRNAs (*36*), its effect on CD4 T cell activation and proliferation is unclear. To address this, we measured protein synthesis rate in activated CD4 T cells 24h post RocA treatment using the puromycin incorporation assay (*37, 38*). RocA-treated cells showed a dose-dependent decrease in puromycin incorporation, suggesting inhibition of protein synthesis. We also found that RocA treatment inhibited cell-proliferation in a dose-dependent manner at sub-nanomolar concentrations (Figures 6A, B). Surprisingly, early T cell activation genes such as IL-2 and CD25 (IL-2Rα) were unaffected both at the mRNA and protein level by RocA, suggesting that impacting early TCR signaling dependent on NFAT is not the mechanism by which RocA inhibits T cell proliferation. Furthermore, this supports the notion that translational blockade through RocA impacts only a subset of mRNAs even in T cells. As expected, when cells were treated with the calcineurin inhibitor Tacrolimus (FK506), T cell proliferation was inhibited, with a corresponding downregulation of both IL-2 and CD25 (IL-2Rα) (Figure 6C). These results suggest that while both RocA and FK506 inhibit CD4 T cell activation and proliferation, they do so through distinct mechanisms. While our finding is different from a previous study suggesting that RocA inhibits NFAT activity in human CD4 T cells (*39*), we propose at higher concentrations (50-100 fold) of RocA could affect transcription regulation. Nevertheless, these data demonstrated that RocA-mediated translation inhibition could suppress CD4 T cell proliferation *ex vivo*.

**Figure 6.**
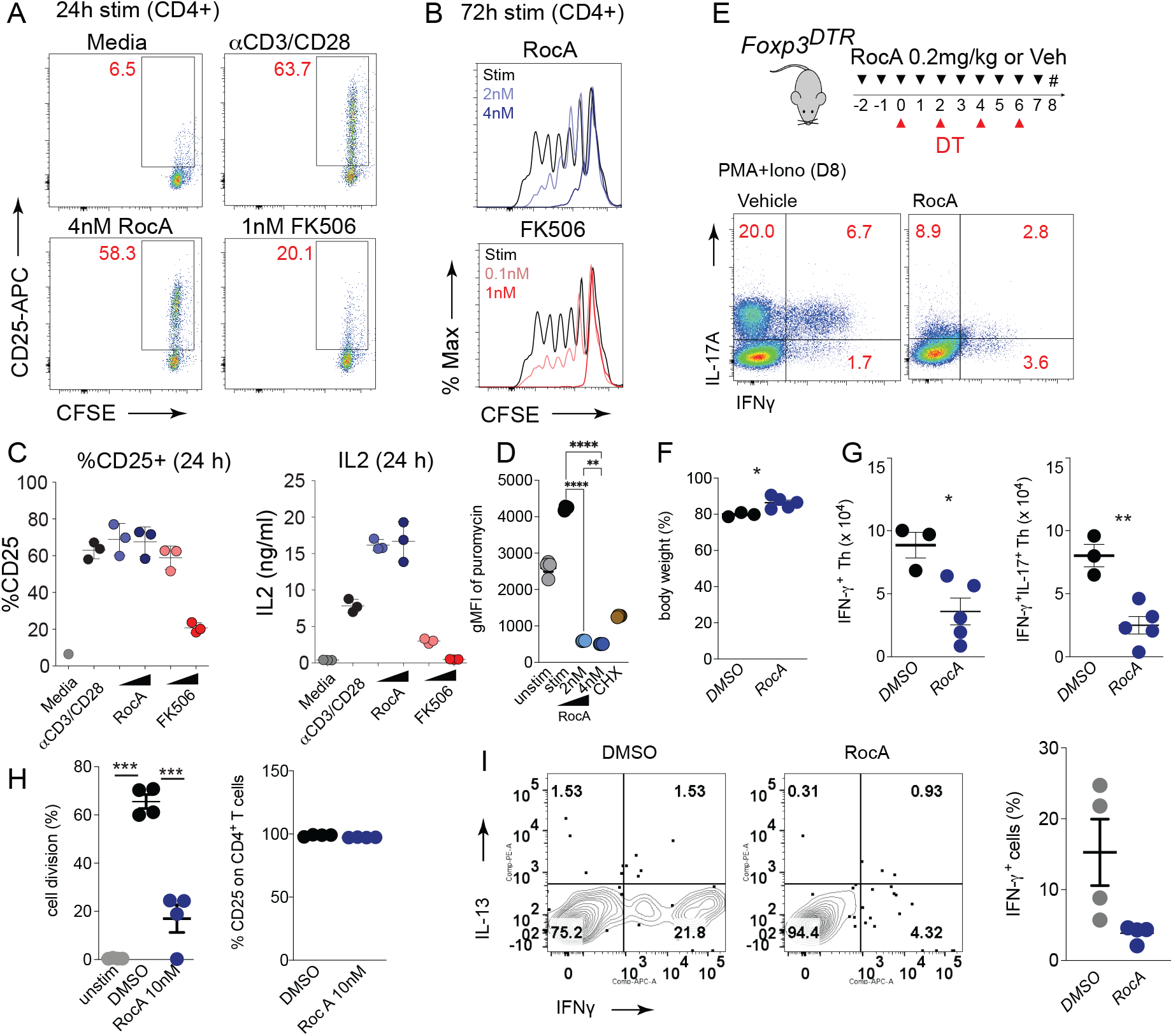
Acute inflammation due to Treg deficiency was reversed by an mRNA translation inhibitor. (A-C) Mouse naïve CD4 T cells were isolated from the spleens of naïve C57BL/6 mice and CFSE labeled. The cells were stimulated with equal number of anti-mouse CD3/CD28-conjugated beads in the presence of the indicated concentration of RocA, FK506, or vehicle for 24h (A, C) or 72h (B). The cells were stained for anti-CD4 (A and C) or anti-CD4 and anti-CD25 (B) for flow cytometric analysis. The data of frequency of CD25 in A was summarized in C. Murine IL-2 was assayed by ELISA from supernatants collected after 24h after stimulation (C). (D) Incorporation of puromycin analyzed by flow cytometry post-RocA treatment. (E-G) Foxp3^DTR^ mice were treated with diphtheria toxin (DT) every other day and 0.2 mg/kg of RocA everyday starting from 2 days before DT treatment, followed by isolation of splenocytes at day 8. The cells were stained with anti-CD4, anti-TCRβ, and anti-CD44 followed by restimulation with PMA and Ionophore for 6h. The cells were stained by anti-IL-17A and anti-IFNγ, followed by flowcytometric analysis (E). The body weight of mice at day 8 (F). The number of Th1 cells (IFNγ^+^CD44^hi^CD4^+^TCR-β^+^) and pathogenic Th17 cells (IFNγ^+^IL-17A^+^CD44^hi^CD4^+^TCR-β^+^) (G). (H) Human naïve CD4 T cells isolated from frozen PBMC, subjected to CFSE labelling. The cells were stimulated with anti-human CD3 and anti-human CD28-conjugated beads in the presence of 10 nM of RocA or vehicle or without the beads indicated as unstim. After 3 days, cells were stained with anti-human CD25 and anti-CD4, subjected to flow cytometry to analyze the frequency of divided cells and CD25+ cells. (I) Human memory Th1 cells isolated as CD4^+^CD25^-^CD45RA^-^CXCR3^+^CCR6^-^ cells from frozen PBMC were incubated in the presence of 10 nM of RocA or vehicle for 2 days, followed by restimulation with PMA and Ionophore for 6h. The cells were surface stained by anti-CD4 and subjected to cytoplasmic staining for IFN γ and IL-13. The frequency of IFNγ^+^ cells is summarized in the dot plots. One-way ANOVA was used for C, D and H; Two-tailed *t*-test (unpaired) was applied for F, G and I. \**P*<0.05, \*\**P*<0.01, \*\*\**P*<0.001. The above experiments were repeated at least twice.

### Acute inflammation due to Treg deficiency is counteracted by the mRNA translation inhibitor Rocaglamide A

We next tested if *in vivo* RocA administration could ameliorate inflammation induced CD4 T cell activation. First, we assessed the effect of RocA on the general health of the mice and immune homeostasis by treating mice every other day for a week with RocA (0.5 and 1mg/kg i.p.) (Figure S6). RocA neither impacted overall weight, or the relative contributions of various cell populations within the CD45^+^ compartment, including CD4 T cell subsets (Figure S6C-E). Thus, short-term *in vivo* treatment with RocA induced no deleterious effects on immune homeostasis and was well tolerated by the mice.

We next asked whether inflammatory responses were limited by RocA treatment in the absence of Tregs. For these studies we used the Foxp3^DTR^ mice, treated with DT to deplete Tregs as described in Figure 1D. To test the ability of RocA to affect acute inflammation, we dosed Foxp3^DTR^ mice with RocA (0.2 mg/kg i.p.) or DMSO (vehicle) on days −1 and −2, and then treated with DT (100 ng i.p.) on days 0, 2, 4, and 6, followed by sacrifice and analysis on day 8 (Figure 6E). A hallmark feature of Treg depletion is a rapid loss of body weight, which is seen in the vehicle-treated mice on day 7, with an average weight loss of 20% of initial body weight (Figure 6F). We found that the RocA-treated mice showed significantly less weight loss, averaging between 85-90% of starting weight, suggesting that RocA treatment was blunting the inflammation caused by Treg depletion. Consistent with this, we found that numbers of splenic CD4 T cells producing IL-17A or IFNγ or both cytokines were significantly reduced in RocA-treated mice (Figure 6G). Especially important was the reduction in IFNγ/IL-17A co-producers, as this subset has been found to be pathogenic in inflammatory settings (*40–43*). These data demonstrate that RocA-mediated direct translation inhibition can ameliorate the inflammation seen in mice lacking functional Tregs, supporting our findings that Treg-mediated immune suppression of CD4 T cells is through active translation control targeting a subset of mRNAs required for protein synthesis itself.

Lastly, we also found suppression of proliferation in primary human T cells treated with RocA and stimulated with anti-CD3/CD28 (Figure 6H). Similar to what was seen in mouse CD4 T cells, there was no difference in the surface expression of CD25 (Figure 6H). We also observed significant suppression of IFNγ producing cells with RocA treatment compared to the vehicle control post PMA/ionomycin stimulation (Figure 6I). Altogether, we show herein a novel mechanism of Treg-mediated suppression of CD4 T cell activation which acts through active mRNA translation control to downregulate the protein dosage of protein synthesis machinery components in activated CD4 T cells. Finally, we show in a physiologically relevant setting that direct inhibition of protein synthesis using a small molecule inhibitor (RocA) can have therapeutic efficacy in alleviating unwanted inflammatory CD4 T cell activation.

## DISCUSSION

Regulatory T cells possess the ability to potently suppress effector CD4 T cell responses and maintain proper immune homeostasis. Several mechanisms have been proposed for Treg-mediated suppression, all predominantly focused on inhibiting aspects of effector T cell proliferation. Despite this emphasis, it is still unclear how target cell gene expression is modulated by Tregs. One key hallmark of CD4 T cell activation is the significant upregulation of global protein synthesis capacity and rate. However, the mechanisms of Tregs mediated suppression of proliferation and activation of Teffs is unclear. While Treg-mediated suppression mechanisms have been heavily studied from the Treg-centered perspective, the molecular events within effector CD4 T cells upon Treg encounter have remained unclear (*10, 44, 45*). To address this, we investigated the effect of Tregs on post-transcriptional gene regulation in Teffs, focusing on protein biosynthesis. We observed that Teffs rapidly increased their global protein synthetic rate following TCR-mediated activation, and that Tregs could suppress this increase in translation at 24 hours of activation. Interestingly, we found Tregs inhibited this process by shutting down the biosynthetic ramp-up required for subsequent cell division, as early as 6h following T cell activation. Similarly, an acute depletion of Tregs *in vivo* led to the robust upregulation of protein synthesis rate in peripheral CD4 T cells. These data suggest that an important aspect of Treg-mediated suppression is inhibition of the ability of Teffs to increase their biomass prior to cell division. The rapid increase in protein synthesis following acute loss of Tregs demonstrates a need for Tregs to hold in check the aberrant increase in protein synthesis of autoreactive T cells in the periphery. This is a highly efficient mode of regulation: inhibiting the ramp-up of biosynthesis that proceeds cell division, thereby stopping proliferation before it has begun.

To uncover the mechanism by which Tregs suppress protein synthesis in Teffs, we developed a novel technique called SPEED to investigate the translatome. We found that, upon activation, Teffs displayed a rapid redistribution of mRNAs from monosome to polysomes, leading to increased translation. This redistribution was independent of acute changes in the transcription of the mRNAs. Most of these mRNAs encoded for proteins important for protein biosynthesis, including ribosomal proteins, RNA-binding proteins, and elongation and splicing factors, suggesting that this process was involved in increasing the biomass prior to cell division. Importantly, we found that activation in the presence of Tregs inhibited the redistribution of these mRNAs, causing them to remain either free or associated with monosomes. As these changes in ribosome occupancy were independent of gene transcription and occurred within 24 hours of stimulation, they would be undetectable by the changes in gene expression and cell division, employing traditional methods to study Treg-mediated suppression. Interestingly, the mRNAs which has higher TE and affected by Tregs were enriched for 5’ UTR terminal oligopyrimidine (TOP) motifs. This motif is enriched in 5’UTR of mRNAs including ribosomal proteins and elongation factors that are core proteins of the translational machinery. This is a motif recognized by the RNA binding protein LARP1 which regulates the association of these mRNAs with stress granules (*46, 47*). The presence of this motif on Treg-affected mRNAs suggests that Tregs may regulate the trafficking of these mRNAs from ribosomes to stress granules, thus controlling their translation.

From the perspective of the requirements for efficient Teff activation and clonal expansion, it is essential that activated Teffs synthesize new proteins and increase their biomass. We observed that this process is specifically regulated by mTORC1 signaling. While the mTOR pathway has been previously implicated to be critical for coordinating cell growth and proliferation in lymphocytes(*11, 48*), we observe that Tregs specifically block mTORC1 signaling required for mRNA translation. We found that activation in the presence of Tregs led to an inhibition of S6 and 4E-BP1/2 phosphorylation, both downstream targets of mTORC1. As mRNA translation is energetically demanding, an mTOR dependent switch to aerobic glycolysis is well known(*49, 50*). Recent study has shown key enzymes in the pathway are translationally regulated important for metabolic reprogramming of activated cell(*14*). These data suggest that translation remodeling occurs earlier than metabolic reprogramming in these cells. We did not detect any mRNAs linked to metabolic processes as we probed for early events of translation remodeling. Similarly, acute depletion of Tregs *in vivo* led to the acute upregulation of protein synthesis rate in peripheral CD4 T cells. This suggests that there is a need for Tregs to hold in check the aberrant increase in protein synthesis of autoreactive T cells in the periphery.

We demonstrated that Treg-derived IL-10 and TGFβ were critical for the ability of Tregs to exhibit translation control in Teffs. Blockade of each individually lead to moderate restoration of translation, while blockade of both cytokines nearly recapitulated translation in the absence of Tregs. Similarly, addition of IL-10 or TGFβ during Teff stimulation (in the absence of Tregs) resulted in a reduction of global translation similar to that seen when Teff cells were stimulated in the presence of Tregs. Thus, IL-10 and TGFβ signaling in Teff cells is involved in Treg-mediated trans-inhibition of protein synthesis. Previous studies have shown divergent effects of IL-10 signaling on mTORC1 activity. In macrophages, IL-10 acts in a paracrine or autocrine fashion to inhibit mTORC1 activation following LPS stimulation(*51, 52*). The outcome was a metabolic reprogramming of the macrophages to promote oxidative phosphorylation. However, in human NK cells, IL-10 regulated metabolic changes that enhanced cellular function (e.g., IFNγ production), and these changes were mediated via activation of mTORC1(*53*). While these studies show opposite effects of IL-10 on cellular activity, they focus on changes in metabolism mediated through regulation of mTOR signaling. However, these studies do not address the role of IL-10 on protein synthesis. Likewise, TGFβ signaling has been shown in a variety of systems to target mTOR, with the outcome dependent on the cell type and the inflammatory context. For example, TGFβ treatment of lung fibroblasts leads to synthesis of collagen through activation of mTORC1 and 4E-BP1/2, the opposite of what we observed in Teff cells(*54*). On the other hand, TGFβ signaling was shown to inhibit mTOR activity in CD8 precursor exhausted cells (Tpex), leading to improved mitochondrial activity during chronic viral infection(*55*). Finally, work from Komai and co-workers (*56*) showed that IL-10 and TGFβ work synergistically to suppress both glycolysis and oxidative phosphorylation via inhibition of mTORC1 in TLR-stimulated B cells. While it is clear from all of these studies that TGFβ and IL-10 can target mTOR for regulation, none of this work addresses the role of these cytokines in regulating translation in CD4 Teffs.

Finally, to determine whether inhibition of translation was an effective method of regulating cellular activation we treated Treg-depleted mice with the mRNA translation inhibitor RocA. We choose RocA as it is a specific inhibitor of mRNA translation, which was well-tolerated *in vivo*. CD4 T cells in mice with acute Treg depletion display an immediate increase in global protein synthesis. Consistent with our hypothesis that inhibition of mRNA translation is an effective means of maintaining tolerance, treatment of these mice with RocA abrogated enhanced protein synthesis and inhibited induction of cytokine expression in Teffs. Taken as a whole, our data show that Tregs suppress Teff activation, through inhibition of mTORC1, leading to suppression of protein translation through sequestration of specific mRNAs from polysomes.

Regardless of the different mechanisms that Tregs may utilize for Teff suppression, a common theme is that changes in intracellular signaling events in Teffs result in inhibition of proliferation. From the perspective of the requirements for efficient Teff activation and clonal expansion, it is essential that activated Teffs synthesize new proteins. We observed that this process is specifically regulated by mTORC1 signaling and Tregs specifically block mTORC1 signaling required for mRNA translation (Figure 7). Our study provides a novel mechanism for the induction and maintenance of peripheral tolerance and identifies Treg derived IL-10 along with TGFβ as critical the regulators of mRNA translation affecting proliferation and activation of effector T-cells.

**Figure 7.**
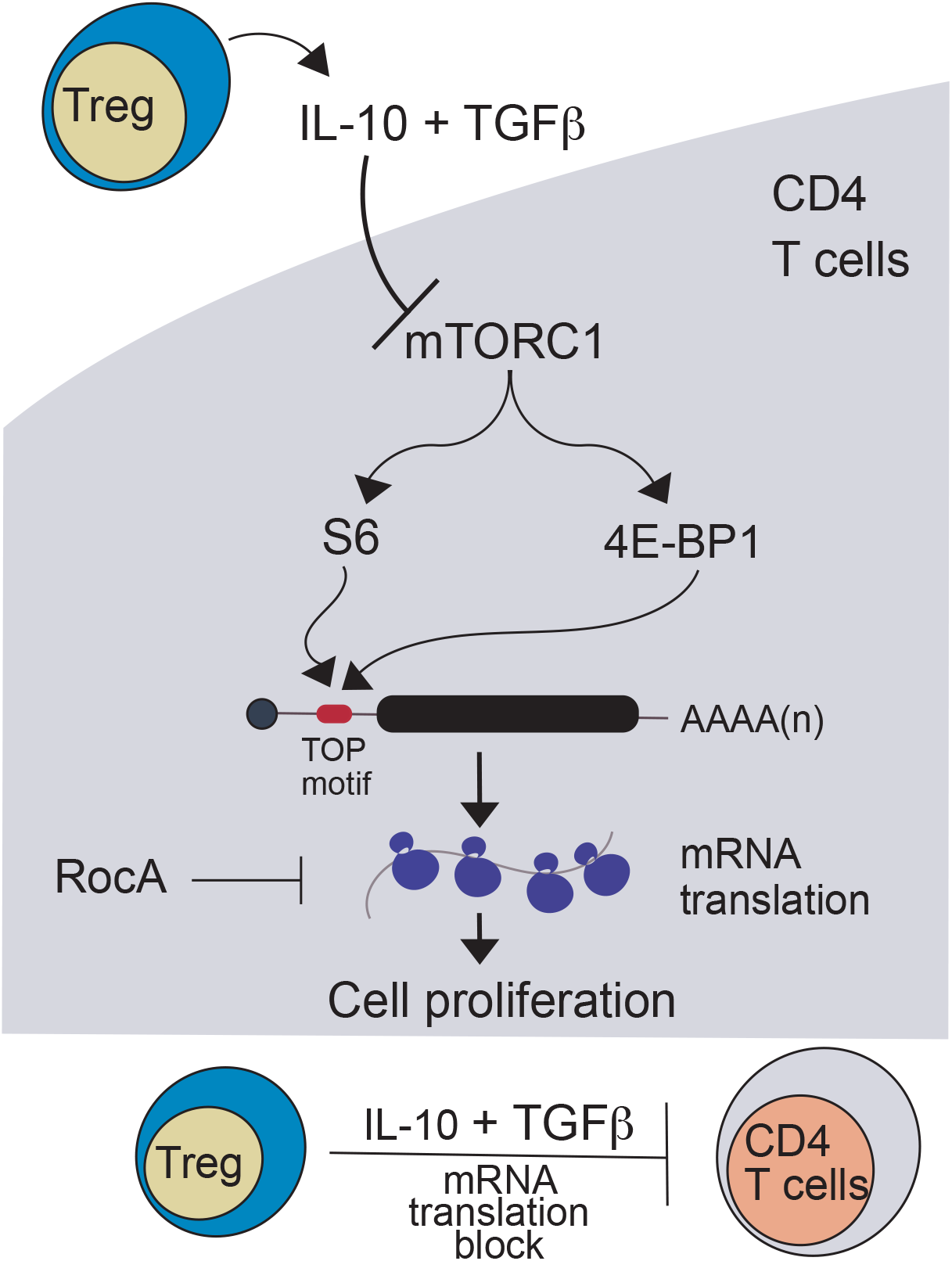
A model depicting Treg mediated disruption of mTORC1signaling that leads to the suppression of protein synthesis of mRNAs enriched for TOP motifs.

## Materials and Methods

### Mice

For Treg-depletion studies, Foxp3^DTR^ mice (B6.129(Cg)-Foxp3*^tm3(DTR/GFP)Ayr^*/J: Stock# 016958 from Jackson Labs) were used. For Treg isolation and expansion *in vivo, Foxp3^YFP-Cre^* mice (B6.129(Cg)-Foxp3*^tm4(YFP/icre)Ayr^*/J: Stock#016959 from Jackson Labs) that have been crossed to B6 Cd45.1 mice (B6.SJL-*Ptprc^a^Pepc^b^*/BoyJ: Stock# 002014 from Jackson Labs) in house were used. For the studies of IL-10 in Treg-mediated suppression of protein synthesis, 7 weeks old female of IL-10RB^-/-^ (B6.129S2-Il10rbtm1Agt/J: Stock# 005027 from Jackson Labs) mice were used. For all the reagent identifiers see Supplementary Table 3.

### In vivo treatments

Foxp3^DTR^ mice were intraperitoneally injected with 100 ng of Diphtheria toxin (DT) at indicated days. For treatment of rocaglamide (RocA), mice were intraperitoneally administrated with indicated concentrations of RocA.

### Murine T cell isolation, culture and suppression assays

Bulk CD4 T cells were isolated from indicated mice and stained with antibodies against CD25, CD44, CD62L, and CD4 except for CD4 T cells from *Foxp3^YFP-Cre^* mice since Foxp3^+^ Tregs were identified using YFP fluorescence from these mice. Purified YFP^-^CD25^-^CD4^+^CD44^low^CD62L^+^ naïve CD4 T cells as CD4 Teffs were sorted using the BD FACS Aria fusion. Unless noted, all isolated T cells were cultured in T cell media (RPMI-1640, 10% Fetal Bovine Serum, 2mM GlutaMAX^tm^-I, 100U/mL Penicillin-Streptomycin, 55μM 2-Mercaptoethanol, 1mM Sodium Pyruvate, 1X Non-Essential Amino Acids, 10mM HEPES). For Treg suppression assays, indicated numbers of tTregs (CD4^+^YFP^+^) and 5 × 10^5^ CD4 Teffs (YFP^-^CD25^-^CD4^+^CD44^low^CD62L^+^ naïve CD4 T cells) were cultured in round-bottom 96-well plate for the indicated times with 5 × 10^5^ anti-CD3/CD28 coated magnetic beads (Teff:beads = 1:1). To control for T cell density, control cultures were cultured with either the same number of total Teffs as in the Treg:Teff co-culture conditions or in some cases, YFP^-^CD25^-^ CD4^+^CD44^low^CD62L^+^CD45.1 ^+^ Teffs sorted from *Foxp3^YFP-Cre^* mice were used. To test proximal TCR signaling, in vitro Treg suppression assay was performed as described with modifications. CD4 Teffs were isolated from Nur77-GFP mice and Tregs were sorted from *Foxp3^YFP-Cre^* mice. Prior to co-culture, CD4 Teffs were further stained with CellTraceViolet to assess proliferative status and ensure proximal TCR signaling assessed by Nur77-GFP was prior to proliferation of these cells. Unstimulated Nur77-GFP CD4 Teffs and Tregs from wild-type C57BL/6 mice served as a negative for GFP signal.

For puromycin incorporation (as a measure of global protein synthesis) puromycin (10 mg/ml) was added for the last 15 minutes of culture, and then the cells were permeabilized and stained with Alexa647-coupled anti-puromycin antibody and analyzed by flow cytometry. Controls included unstimulated Teffs also labeled with puromycin, and stimulated Teff given CHX at the same time as puromycin (CHX results in a complete translation blockade, so this controls for non-specific uptake of puromycin). For analysis of Treg-mediated suppression we have presented the puromycin incorporation data as % suppression, relative to the puromycin incorporation in the Teff alone condition (which is put at 0% inhibition). In this calculation, 100% of suppression means absolute inhibition of puromycin incorporation. For analysis of mTORC signaling, we have presented the geometric mean fluorescence intensity (gMFI) data as normalized gMFI, relative to the gMFI data in the Teff alone condition (which is put at 1).

### Ex-vivo human Treg suppression assay

We used in vitro-expanded Tregs from a single donor, and Teff (defined as CD4+CD45RA+) from 5 individual healthy donors for Treg suppression assay. Established protocols were used for Treg expansion (*57, 58*) and Teff (CD4^+^CD45RA^+^) cells were purified PBMC using the naïve human T cell isolation kit from Miltenyi. To determine the effect of Tregs on overall translation in Teff we cultured Teff (10^4^ cells) in absence or presence of Tregs (1:4 Treg:Teff) for 24 hours with anti-CD3/CD28 beads (at a ratio of 28:1 [Teff:beads]). Puromycin (10 μg/ml) was added for the last 15 minutes of culture, and then the cells were permeabilized and stained with Alexa647-coupled anti-puromycin antibody and analyzed by flow cytometry. Controls included unstimulated Teffs also labeled with puromycin, and stimulated Teff given CHX at the same time as puromycin (CHX results in a complete translation blockade, so this controls for non-specific uptake of puromycin).

### Low input sucrose gradient polysome fractionation

Sucrose gradients (15-60%) were prepared in SW55Ti rotor-compatible Ultra-Clear ultracentrifuge tubes (Beckman Coulter, 344057). Briefly, a 2M (68.5%) sucrose solution in nuclease-free water and a 10X sucrose buffer (250mM Tris-HCl pH 7.5, 1.5M KCl, 150mM MgCl2, 10mM DTT, 1mg/ml CHX, Complete protease inhibitor EDTA-free (2 Tablets per 50ml), 20U/ml SUPERaseIn RNAse Inhibitor) was prepared. Using these two solutions, sucrose solutions with 1X sucrose buffer and different concentrations (60%, 45%, 30%, 15%) were prepared. Each solution was added from the bottom of the tubes in the following order and quantity (60%: 750μl, 45%: 1.5ml, 30%: 1.5ml, 15%: 750μl). For each addition, tubes were kept in the −80C for at least 15min to freeze the sucrose solutions before adding the next sucrose solution. All tubes were sealed with parafilm and kept at −80C forever. Tubes were allowed to thaw at 4C for 12-16h before the fractionation. Samples were prepared in PLB buffer similar to the Ribo-IP method and total RNA was quantified using RiboGreen in the low-range assay (1ng/mL – 50ng/mL). All samples were adjusted with PLB to at least 500ng of total RNA and layered carefully on top of thawed sucrose gradient tubes. Tubes were ultracentrifuged at 35,000rpm in a SW55Ti rotor using the L8-70M ultracentrifuge (acceleration: default, deceleration: 0). Separated lysates were fractionated from top to bottom with an Auto-Densi Flow fractionator (Labconco) with continuous reading of absorbance at 254nm (A254nm) using a UA-6 UV/VIS detector (Teledyne Isco). A total of 16-18 fractions (250-300μl) were fractionated with the Foxy R1 fractionator in 2ml Eppendorf tubes and kept on ice. For digital conversion of the A254nm signal, we attached the LabQuest Mini data-collection interface (Vernier, LQ-MINI) to the UA-6 detector. For RNA extraction, we used 3 volumes of Trizol LS to each sucrose fractions and vortexed vigorously. Every fraction was spiked-in with *in vitro* transcribed firefly Luciferase RNA (uncapped) to assess RNA extraction efficiency between fractions. Samples were either kept in −80C at this point or proceeded with standard Trizol-mediated RNA extraction protocol or the Direct-zol 96 kit (Zymo Research, R2054) was used following the on-column DNA digestion protocol. Total RNA quantity from each fraction was measured using RiboGreen in the low-range assay to generate ribosome traces. The Bioanalyzer 2100 RNA pico kit was also used to assess the starting point of the intact 80S monosome peak (containing both 28S and 18S rRNA bands). Equal volumes of RNA from fractions corresponding to polysomes (3 fractions after the 80S monosome peak) were pooled and digested with RQ1 DNase (Promega) and cleaned-up using RNA Clean&Concentrator-5 kit (Zymo Research, R1013). Finally, RNA quantity (RiboGreen) and RNA quality (Bioanalyzer) were measured again before proceeding with cDNA libraries construction using the SMARTseq v4 Ultra-Low Input kit (Clontech).

### RNA-seq bioinformatics methods

Reads were aligned by STAR (*59*) 2.4.2a to GRCm38.91, and genes that were not expressed in at least 20% of samples were filtered out. The reads were TPM normalized and modeled using the software LIMMA^(*60*)^. The LIMMA model included a variable for each of the four mice and categorized samples by fractionation (input, polysomal, or sub-polysomal), whether they were stimulated, and their treatment (HHT, T-regulatory cells, or none). Combinations of those conditions resulted in 12 categories total. Translation efficiency was computed as the log fold-change in TPM between the polysomal and sub-polysomal fraction per condition, both at the individual mouse and aggregate levels. Differential translation efficiency was computed in LIMMA as differences between translation efficiencies between conditions. Gene set enrichment statistics were computed from LIMMA-computed log fold-changes and the fgsea (*61*) package. Motif enrichment analysis was performed on the 131 genes whose TE increased in stimulated cells relative to unstimulated cells at 5% FDR and also decreased in TE in Treg-exposed stimulated cells relative to the unstimulated cells. The 5’ untranslated regions for those 131 genes were queried using biomart, which returned sequences for 127 of the genes(*62*). The MEME software tool was used to identify de novo a motif enriched in these sequences (considering both the positive and negative strand)(*63*). The SEA software tool was used to confirm the enrichment of this motif on the strand relevant for mRNA binding(*64*).

### Antibody neutralization assays

Tregs were sorted from CD45.1 *Foxp3^YFP-Cre^* mice as CD4^+^YFP^+^ cells, subjected to labeled with CFSE (Thermo). Indicated number of Tregs were stimulated with anti-CD3 and anti-CD28 antibody-conjugated beads for overnight prior to suppression assay. Naïve CD4 T cells were isolated from WT mice or IL-10Rβ-deficient mice using Mojo sort (biolegend), followed by staining with cell trace blue (Thermo). Indicated number of naïve CD4 T cells was added into the stimulated Treg cells for suppression assay. In some experiments, those naïve CD4 T cells were pre-incubated with 10 mg/mL of neutralizing antibody for IL-2 (JSE6-1A12), IL-2R (3C7; Biolegend), TGFβ (1D11; BioXcell) and/or IL-10R (1B1.3A; BioXcell) for 30 min before adding Tregs and stimulation beads.

## Acknowledgements

This project was funded by National Institutes of Health grants R21AI143227 (SZ & RS), 5T32AI106677-07 (T32) and 1F32AI145283-01A1 (F32) training grant to L.S. We thank the members of the Savan and Ziegler labs for advice and input. We also thank Daniel J Campbell for his advice, Pamela J Fink, Nandan Gokhale and Stephen Anderson for their critical reading and comments.

## Supplementary Figures

**Supplementary Figure 1.**
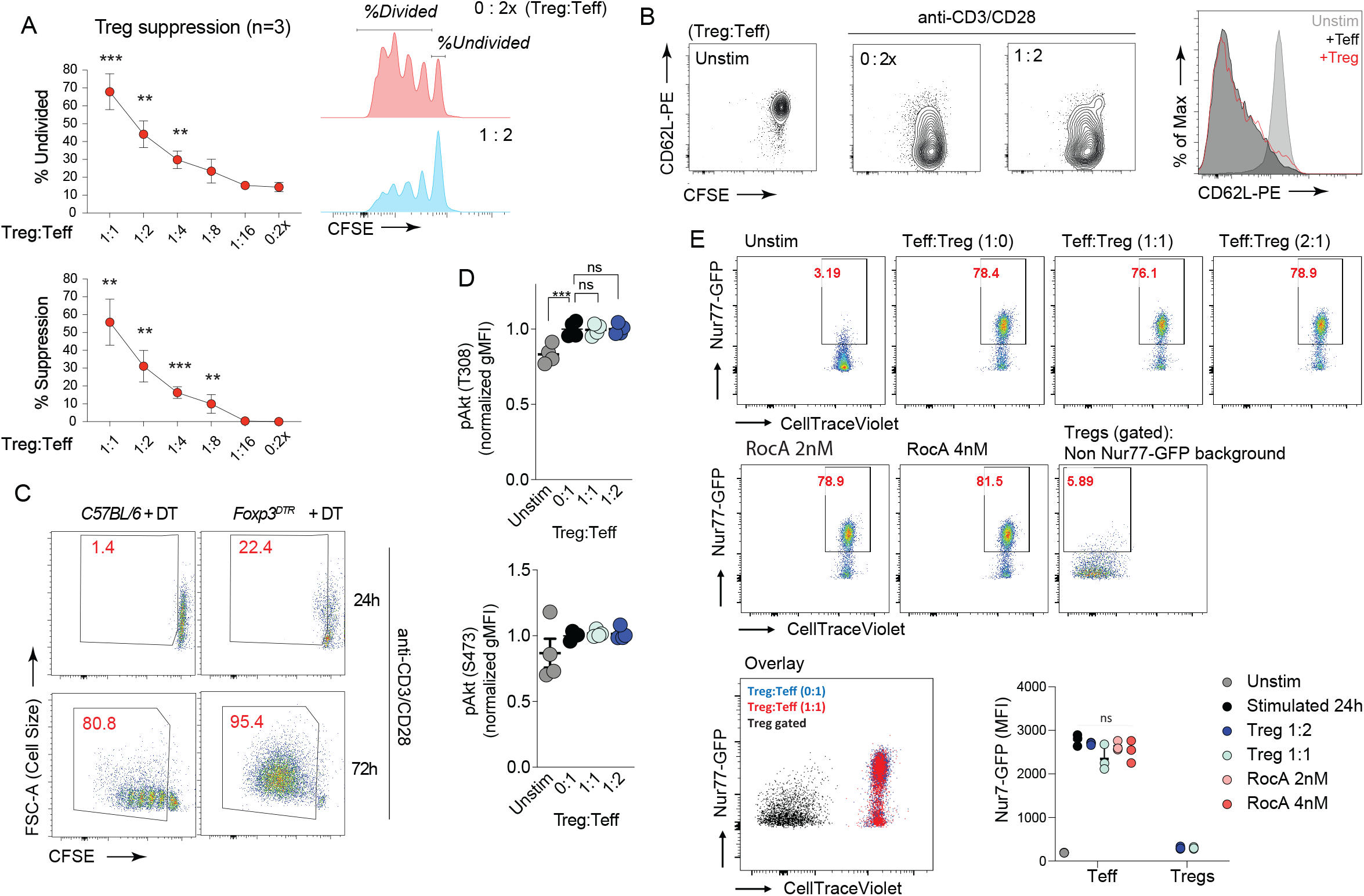
(A) *In vitro* Treg suppression assay using different doses of purified Foxp3-YFP+ Tregs and co-culturing with CFSE-labeled YFP^-^CD25^-^CD4^+^CD44^low^CD62L^+^ naïve CD4 T cells as CD4 Teffs. Data is presented in two ways. Numerating the undivided CFSE peak to show percent undivided cells. Percent suppression was calculated by measuring cells that have divided at least once. (B) CD62L expression was measured in unstimulated and stimulated CD4 Teffs with or without Tregs at 24h post stimulation. (C) Bulk CD4 T cells were purified on day 2 from control C57/B6 mice and Foxp3^DTR^ mice treated with DT for two consecutive days. Cells were labeled with CFSE and activated using anti-CD3/CD28 beads. Cell size (forward scatter) and proliferation (CFSE dilution) was measured at 72h post stimulation. (D) CD4 Teffs stimulated in the absence and presence of Tregs were identified by congenic markers and intracellular signaling molecules were assessed by phospho-flow cytometry. (E) CD4 Teffs from Nur77-GFP mice labeled with CellTraceViolet were either activated with anti-CD3/CD28 beads alone or with titrating amounts of sorted Tregs. Some conditions received indicated concentrations of RocA for eIF4A inhibition. Proximal TCR signaling was assessed using GFP in CD4 Teffs by gating on CellTraceViolet^+^ population. Each datapoint represents a biological replicate (CD4 Teffs from individual Nur77-GFP mice).

**Supplementary Figure 2.**
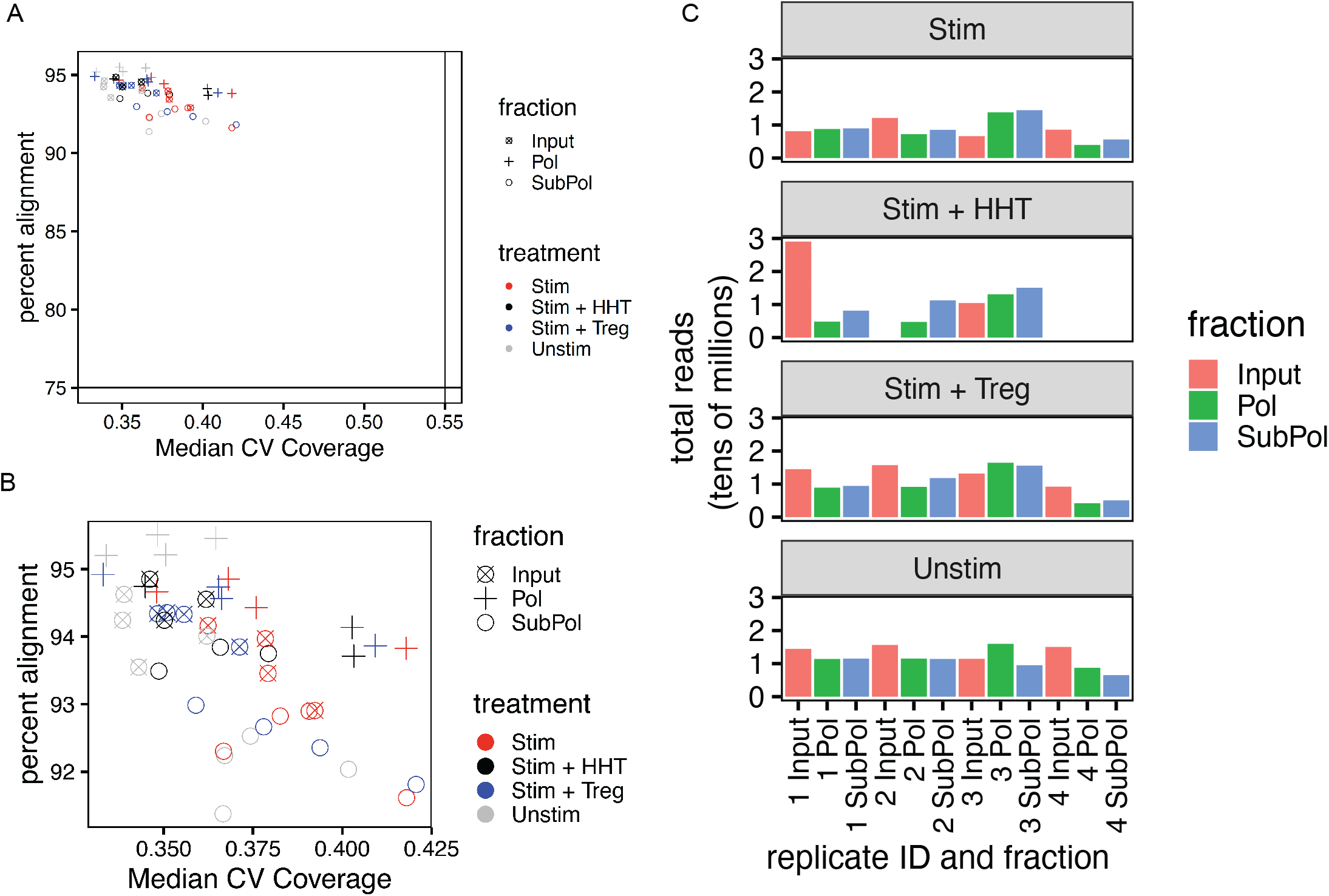
(A) Scatterplot for each sample of quality control metrics percent alignment and median covariance of coverage, colored by fraction and treatment. Percent alignment refers to the percent of reads that can be aligned to the reference genome. Median covariance of coverage refers to the median coefficient of variation (mean/standard deviation) of gene sequence coverage for the 1000 most highly expressed transcripts. All samples pass quality control thresholds of 75% percent alignment and 0.55 median covariance of coverage. (B) Same as panel A, zoomed into the region of the graph where the samples reside. (C) Bar graph of the number of millions of reads sequenced from each sample.

**Supplementary Figure 3.**
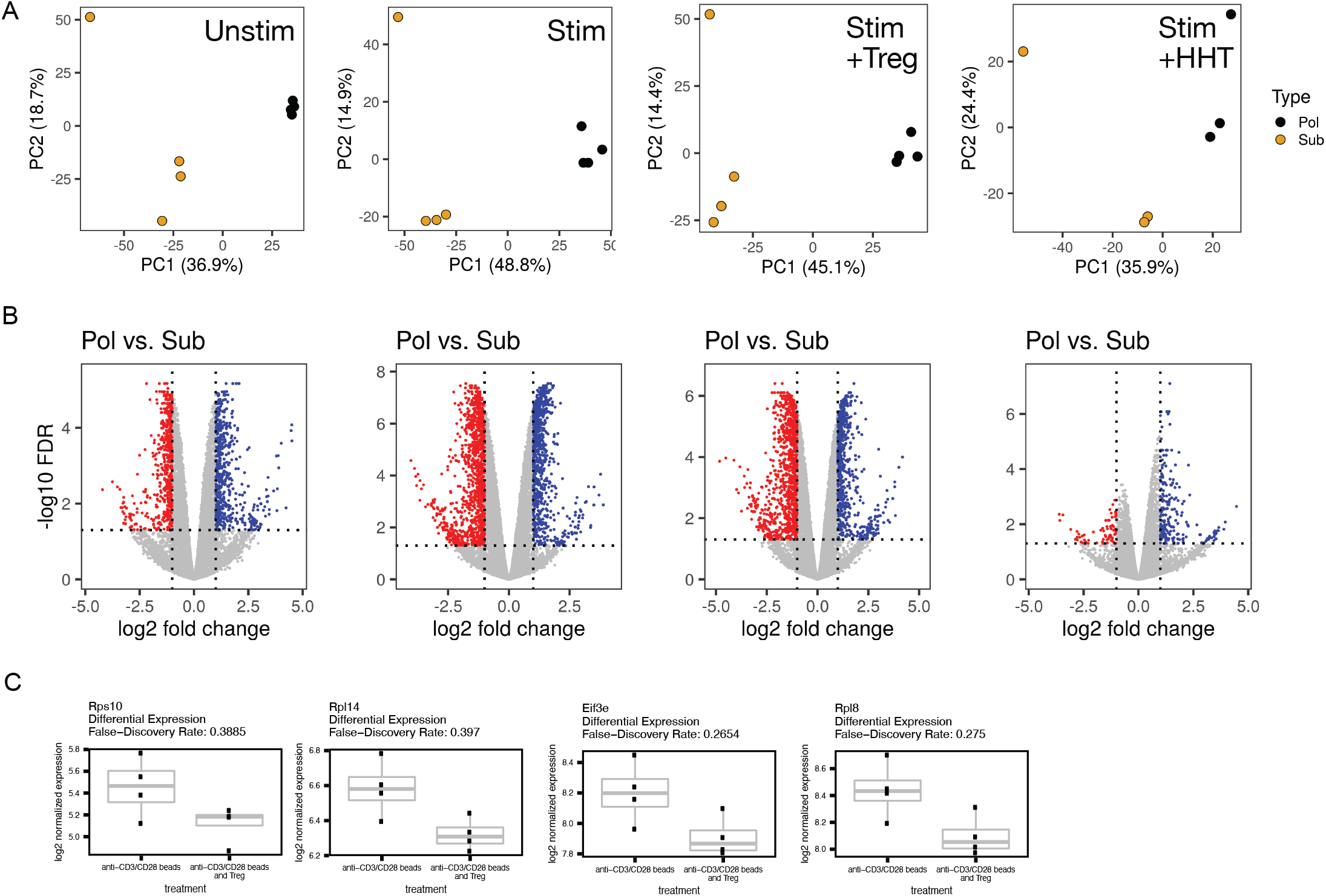
(A) Scatterplots showing principal components 1 and 2 for each sample across the four treatment conditions and two fractions. Principal components were computed separately for each treatment condition. (B) Scatterplots showing translation efficiency (log_2_ fold-change of gene expression between the polysomal fraction and the subpolysomal fraction) and −log_10_ false-discovery rate for each stimulation condition. Translation efficiency false-discovery was computed using the software package LIMMA, and the false-discovery rate refers to Benjamini-Hochberg-corrected p-values for the hypotheses for each gene that the log_2_ fold-change between the fractions does not equal 0. Genes are colored red if their log_2_ translation efficiencies are less than −1 and false-discovery rates are less than or equal to 5%. Genes are colored blue if their log_2_ translation efficiencies are greater than 1 and false-discovery rates are greater than or equal to 5%. (C) Boxplots show the expression of a few translation-associated genes in the input fractions of the bead stimulated and bead stimulated, Treg exposed samples. The false-discovery rates reported are for differential expression between the bead stimulated and bead stimulated and Treg exposed samples, multiple-test corrected across all expressed genes.

**Supplementary Figure 4.**
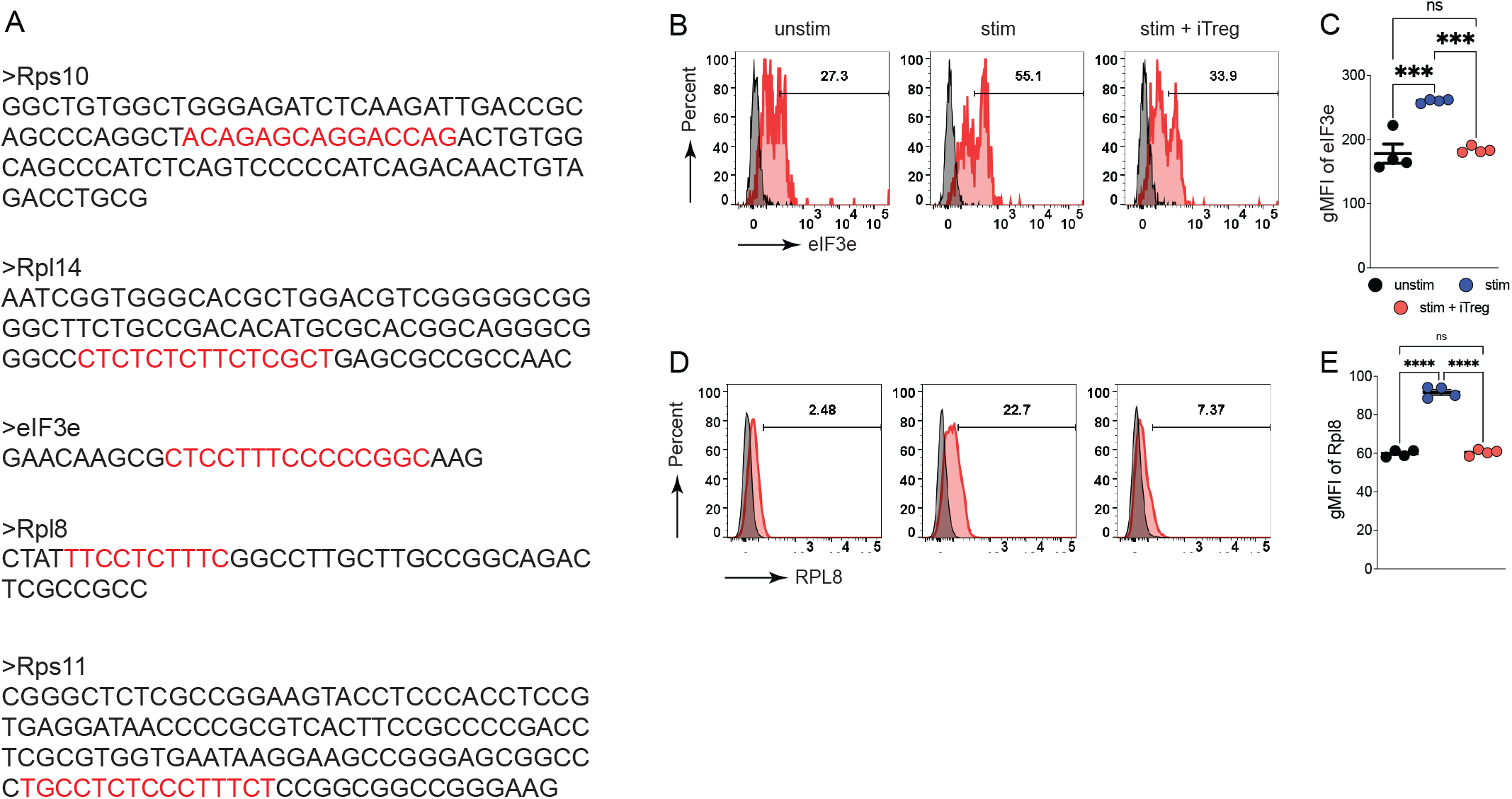
(A)Terminal oligopyrimidine tracts (TOP) motif is highlighted in red for the genes used in Fig 4. (B and D) Purified naive CD4 T cells were harvested with or without equal number of anti-CD3/CD28 beads in the presence or absence of equal number of iTregs for 6h. Histogram plots shows the protein expression levels of RPL8 and eIF3e, analyzed by flowcytometry. (C and E) Summary of the gMFI of the replicates.

**Supplementary Figure 5.**
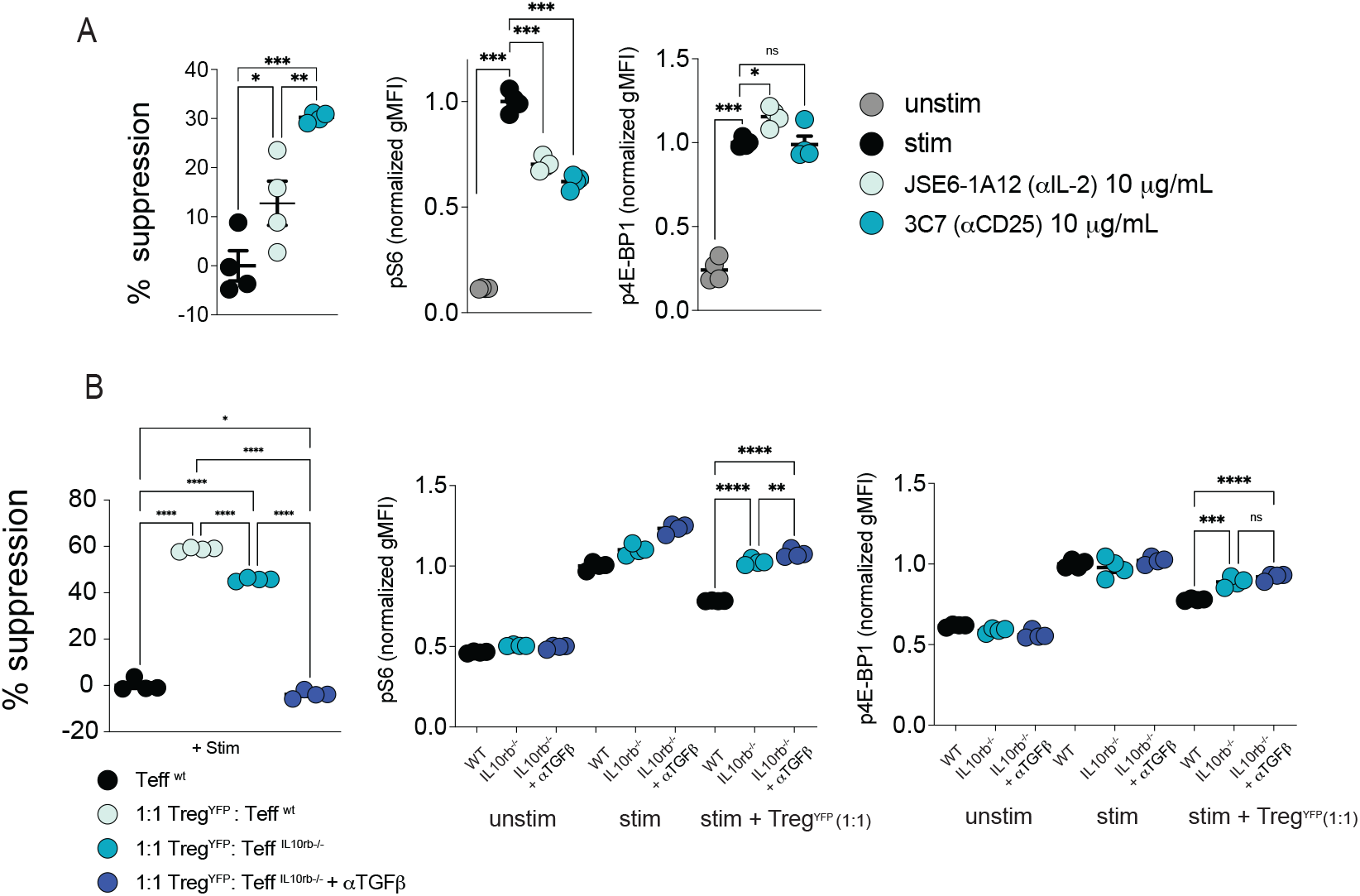
Purified naïve CD4 T cells as CD4 Teffs were harvested with or without equal number of anti-CD3/CD28 beads in the presence or absence of equal number of Tregs for 24h, subjected for analyzing puromycin incorporation and phosphorylation of S6 and 4E-BP1, either alone or with anti-IL-2 or anti-IL-2Ra antibodies (A). (B) CD4 Teffs were isolated from WT mice (black circle) and IL-10Rβ-deficient mice (light blue, cyan, and deep blue circles) were cocultured with or without equal number of anti-CD3/CD28 beads in the presence or absence of equal number of Tregs for 24h and incubated with or without neutralizing αTGFβ, and analyzed for puromycin incorporation, phosphorylation of S6 and 4E-BP1.

**Supplementary Figure 6.**
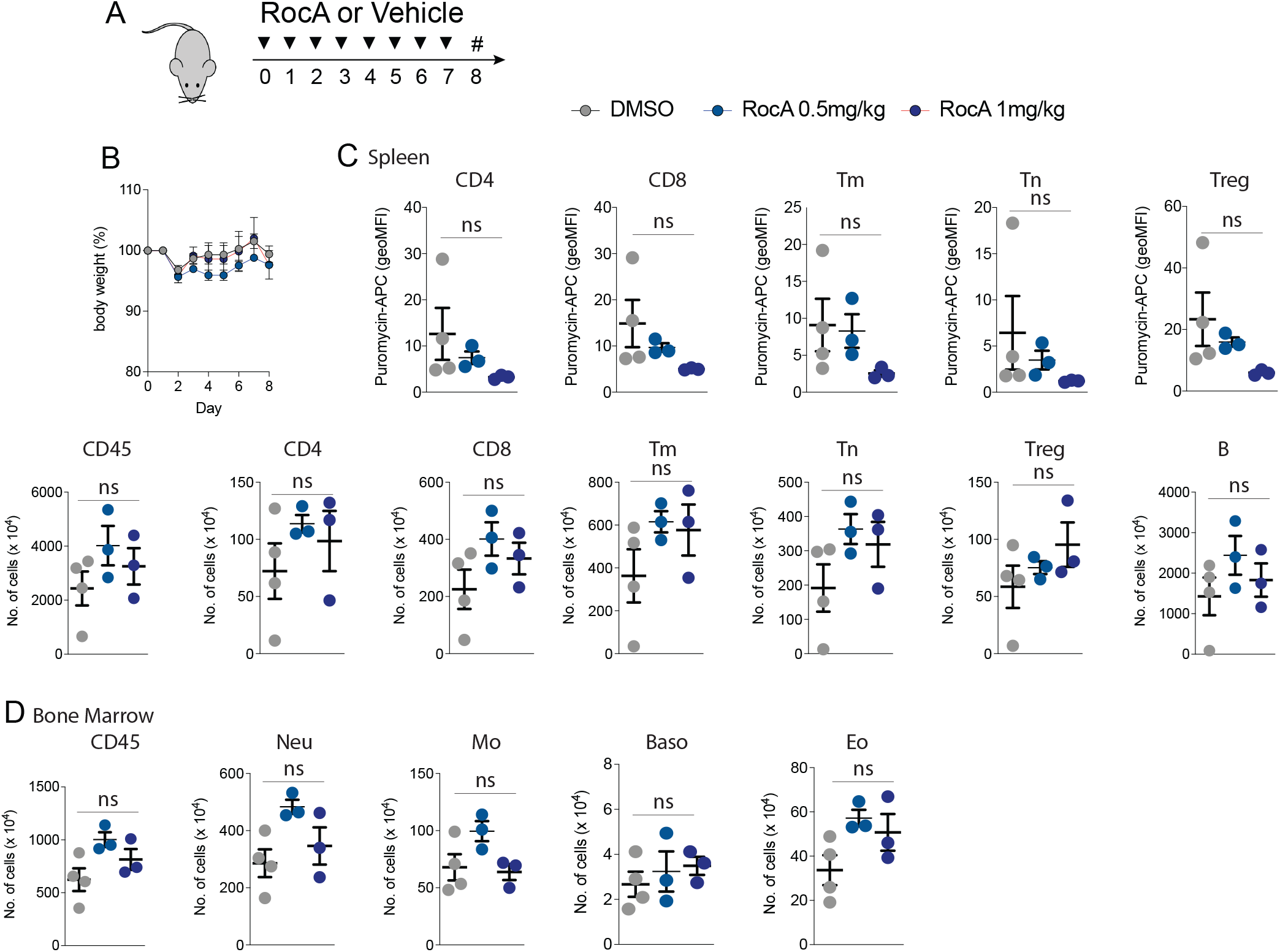
(A) Mice were treated with 0.5 or 1 mg/kg of RocA or vehicle every day for 7 days, followed by isolation of bone marrow and spleen at day 8. Total body weight was monitored everyday post treatment. (B) Incorporation of puromycin analyzed by flowcytometry post-RocA treatment. (B-D) Splenocytes and bone marrow cells and were stained for neutrophils (Ly6G+), eosinophils (SiglecF+CD11c-), basophils (ckit-CD200R3+FceRI+), Monocyte (CD11b+Ly6c+), naïve CD4+ T (CD4+CD3+CD44lowFoxp3-), memory CD4+ T (CD4+CD3+CD44hiFoxp3-), Treg (CD4+CD3+Foxp3+), CD8 T(CD8+CD3+) and B (CD19+) cells in indicated fractions, subjected to flow cytometry.

Supplementary Table 1. List of differentially expressed genes using SPEED technique. These data are presented in Figure 2.

Supplement Table 1-Speed gene list tabs:

*Stim_v_unstim_TE_v_mRNA_input: genes that have differential expression at 5% in the input mRNA in stim vs. unstim, but do not have differential TE in stim m vs. unstim Stim_v_unstim_TE_down: genes that decrease in TE in Stim vs. Unstim at 5% FDR Stim_v_unstim_TE_up: genes that increase in TE in Stim vs. Unstim at 5% FDR Stim_v_unstim_TE_mRNA_both: genes that change in both TE and mRNA expression in the same direction in stim vs. unstim at 5% FDR*

*Stim_v_unstim_TE_mRNA_neither: genes that do not change in TE nor mRNA expression in stim vs. unstim at 5% FDR*

*Stim_v_unstim_TE_v_mRNA_opposite: genes that change in both TE and mRNA expression in stim vs. unstim, but in the opposite direction, at 5% FDR*

*Stim_v_unstim_TE_v_mRNA_TE: genes that change in TE but not mRNA expression in stim vs. unstim at 5% FDR*

*Stim_and_treg_TEs_by_donor: differential log TE in Treg vs. Stim for particular genes, separately for each donor. \**

Supplementary material 2A-E.

*A. TE_StimVNostim_131_DE.csv*

*B. MEME_top_motif_gene_list.txt*

*C. MEME_127_motif_output.txt*

*D. Sea.tsv*

*E. SEA_TOPMotifSequences.tsv*

**Supplementary Table 3.**
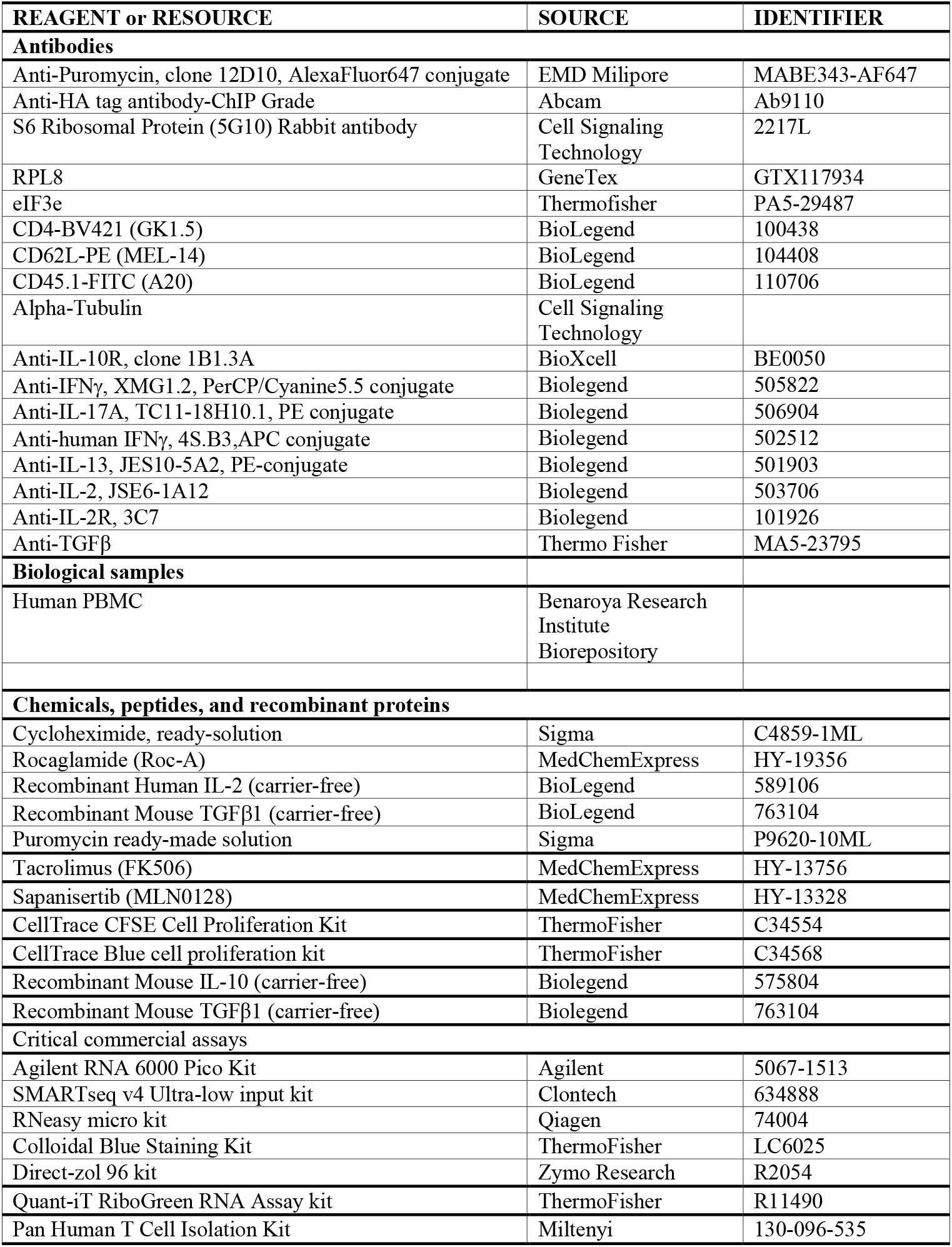
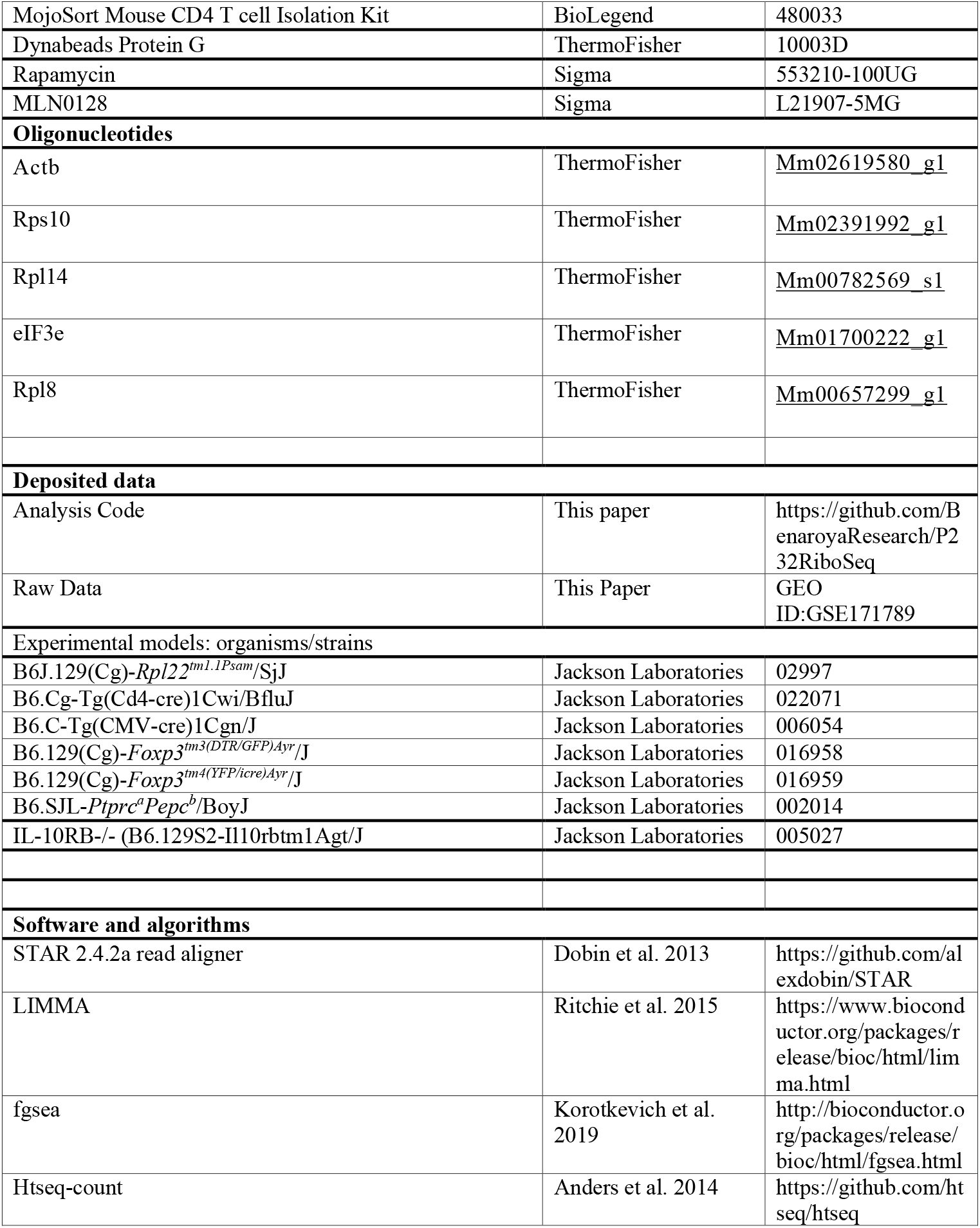
Reagent identifiers. RNAseq dataset identifiers, and computational tools link.

